# Convergent alterations in the tumor microenvironment of MYC-driven human and murine prostate cancer

**DOI:** 10.1101/2023.09.07.553268

**Authors:** Mindy K Graham, Rulin Wang, Roshan Chikarmane, Bulouere Wodu, Ajay Vaghasia, Anuj Gupta, Qizhi Zheng, Jessica Hicks, Polina Sysa-Shah, Xin Pan, Nicole Castagna, Jianyong Liu, Jennifer Meyers, Alyza Skaist, Yan Zhang, Kornel Schuebel, Brian W Simons, Charles J. Bieberich, William G Nelson, Shawn E. Lupold, Theodore L DeWeese, Angelo M De Marzo, Srinivasan Yegnasubramanian

**Author notes:** These authors contributed equally (Mindy K Graham, Rulin Wang). Correspondence: S.Y.

## Abstract

The tissue microenvironment in prostate cancer is profoundly altered. While such alterations have been implicated in driving prostate cancer initiation and progression to aggressive disease, how prostate cancer cells and their precursors mediate those changes is unclear, in part due to the inability to longitudinally study the disease evolution in human tissues. To overcome this limitation, we performed extensive single-cell RNA-sequencing (scRNA-seq) and rigorous molecular pathology of the comparative biology between human prostate cancer and key time points in the disease evolution of a genetically engineered mouse model (GEMM) of prostate cancer. Our studies of human tissues, with validation in a large external data set, revealed that cancer cell-intrinsic activation of MYC signaling was the top up-regulated pathway in human cancers, representing a common denominator across the well-known molecular and pathological heterogeneity of human prostate cancer. Likewise, numerous non-malignant cell states in the tumor microenvironment (TME), including non-cancerous epithelial, immune, and fibroblast cell compartments, were conserved across individuals, raising the possibility that these cell types may be a sequelae of the convergent MYC activation in the cancer cells. To test this hypothesis, we employed a GEMM of prostate epithelial cell-specific MYC activation in two mouse strains. Cell communication network and pathway analyses suggested that MYC oncogene-expressing neoplastic cells, directly and indirectly, reprogrammed the TME during carcinogenesis, leading to the emergence of cascading cell state alterations in neighboring epithelial, immune, and fibroblast cell types that paralleled key findings in human prostate cancer. Importantly, among these changes, the progression from a precursor-enriched to invasive-cancer-enriched state was accompanied by a cell-intrinsic switch from pro-immunogenic to immunosuppressive transcriptional programs with coinciding enrichment of immunosuppressive myeloid and Treg cells in the immune microenvironment. These findings implicate activation of MYC signaling in reshaping convergent aspects of the TME of prostate cancer as a common denominator across the otherwise well-documented molecular heterogeneity of human prostate cancer.

## INTRODUCTION

Like many human cancers, prostate cancer is a highly heterogeneous disease with respect to clinical, pathological, and molecular features. In particular, the ever-growing body of genomic and molecular studies of human prostate cancer has revealed numerous and heterogeneous molecular subtypes and their associations with clinicopathological features ^1–3^.

Despite this heterogeneity, the early stages of prostate cancer development appear to have some convergent patterns, involving transitions from benign glands, to precursor lesions called prostatic intraepithelial neoplasia (PIN), to invasive carcinoma^4, 5^. Luminal epithelial cells in PIN lesions exist within the confines of a normal glandular architecture but display molecular and cellular alterations characteristic of invasive carcinoma cells, albeit with somewhat lower rates. Such truncal alterations, which appear to be early convergent patterns across the known heterogeneity of prostate cancer, include nucleolar enlargement, MYC over-expression, telomere shortening, DNA hypermethylation-mediated silencing of *GSTP1* and other tumor suppressor/caretaker genes, and formation of oncogenic Ets fusions ^6–17^. Furthermore, widespread inflammatory lesions in the prostate, termed proliferative inflammatory atrophy (PIA), are thought to be a hotbed from which PIN and invasive cancer can emerge ^18, 19^. PIA is thought to represent a state of inflammatory injury and regeneration in which proliferative and atrophic epithelial cells can take on an intermediate cell phenotype with both luminal and basal cell characteristics. Consistent with a model implicating PIA as an early step in prostate carcinogenesis, these lesions can be seen to merge directly with PIN and, less frequently, with invasive carcinoma ^20, 21^. Additionally, a subset of PIA cells display some of the convergent molecular patterns seen in PIN and invasive carcinoma lesions, with a low frequency of PIA cells showing MYC over-expression, partial promoter hypermethylation and silencing of GSTP1, and formation of Ets fusions ^14, 18, 22–25^.

The convergent transitions in the luminal epithelial cells during the early development of prostate cancer are accompanied by multiple convergent patterns of alterations in the surrounding microenvironment. The atrophic/intermediate cells observed in PIA lesions generally occur in the context of: i) widespread chronic inflammation composed of myeloid and lymphoid cells, and ii) a reactive stroma with morphologically altered fibroblast and smooth muscle cell compartments that are infiltrated with chronic inflammatory cells. Remarkably, as these PIA regions transition to PIN and invasive cancer lesions, there is a profoundly reduced inflammation and immune cell presence, consistent with the characterization of prostate cancer as “immunologically cold” ^26–28^. Furthermore, basal cells in PIN lesions are relatively depleted compared to benign glands and are completely lost in invasive carcinoma lesions ^6, 29^. Additionally, the fibroblast, smooth muscle, vascular, and other stromal compartments are also thought to be significantly altered in the context of prostatic neoplasia ^30^. However, the precise nature of the epithelial, stromal, and immune cell alterations during the early transitions in prostate carcinogenesis and how the alterations in the neoplastic cells may mediate these changes is poorly understood. This is in part because of a lack of studies focusing on the early stages of prostate cancer development at single-cell resolution and the inability to longitudinally follow these early stages of disease evolution in human tissues.

Here, we have used extensive scRNA-seq, and rigorous molecular pathology approaches to study the comparative biology between human prostate cancer tissues and landmark time points in the disease evolution of a GEMM of murine prostate cancer to define alterations in the transcriptional programs and cellular states during early prostate cancer development. Consistent with the idea that MYC activation in cancer cells can drive changes to the TME ^31^, we show that activation of MYC signaling is a common denominator across the well-known heterogeneity of prostate cancer and that its downstream sequelae mediate a chain of cell network interactions responsible for reprogramming the TME of prostate cancer. Furthermore, we show that prostate-specific activation of MYC signaling in the mouse initially creates a pro-inflammatory state, with many parallels to that seen in human PIA and early PIN lesions, with subsequent transition to an immunosuppressive state during transition to invasive carcinoma, reminiscent of the immunologically cold microenvironment of human invasive prostate cancer.

## RESULTS

### A single-cell transcriptional atlas of localized primary prostate cancer tissues from ten subjects

To develop a detailed atlas of cell states and transcriptional programs in the TME of early prostate cancer, we used scRNA-seq to examine prostate tissues collected from 10 subjects with localized, treatment-naïve prostate cancer that underwent radical prostatectomy. For each subject, tissues were collected from each zone of the prostate (peripheral, central, and transition), as well as from areas with visible tumors (Figure 1A). Cancer-enriched and benign-enriched tissue samples from the peripheral zone (Figure 1B) were used for single-cell RNA-seq assessments, yielding >110,000 cells with high-quality genome-scale gene expression measurements (Figure 1C, Table S1). Dimensionality reduction and cluster analysis revealed the spectrum of cell types present in the prostate TME (Figure 1C-D, Table S2), including luminal and basal epithelial cells, endothelial cells, pericytes, smooth muscle cells, fibroblasts, macrophages, mast cells, T cells, and B cells. Interestingly, in a subset of luminal epithelial cells derived from cancer-enriched tissue samples, we observed a striking subject-specific clustering (Figure 1C), consistent with the known inter-individual tumor heterogeneity in prostate cancer ^1, 32^. However, for the majority of other cell types, there was no subject-specific clustering or skewing; although some clusters, such as basal cells, fibroblasts, and macrophages, did appear to show some skewing in the cancer- vs benign-enriched tissues. These results raised the possibility that although there is significant inter-individual heterogeneity in cancer epithelial cells, there may be cancer-specific convergent alterations in other cells in the TME, evident as less inter-individual heterogeneity. We explored this hypothesis in significant detail in subsequent studies and analyses.

**Figure 1.**
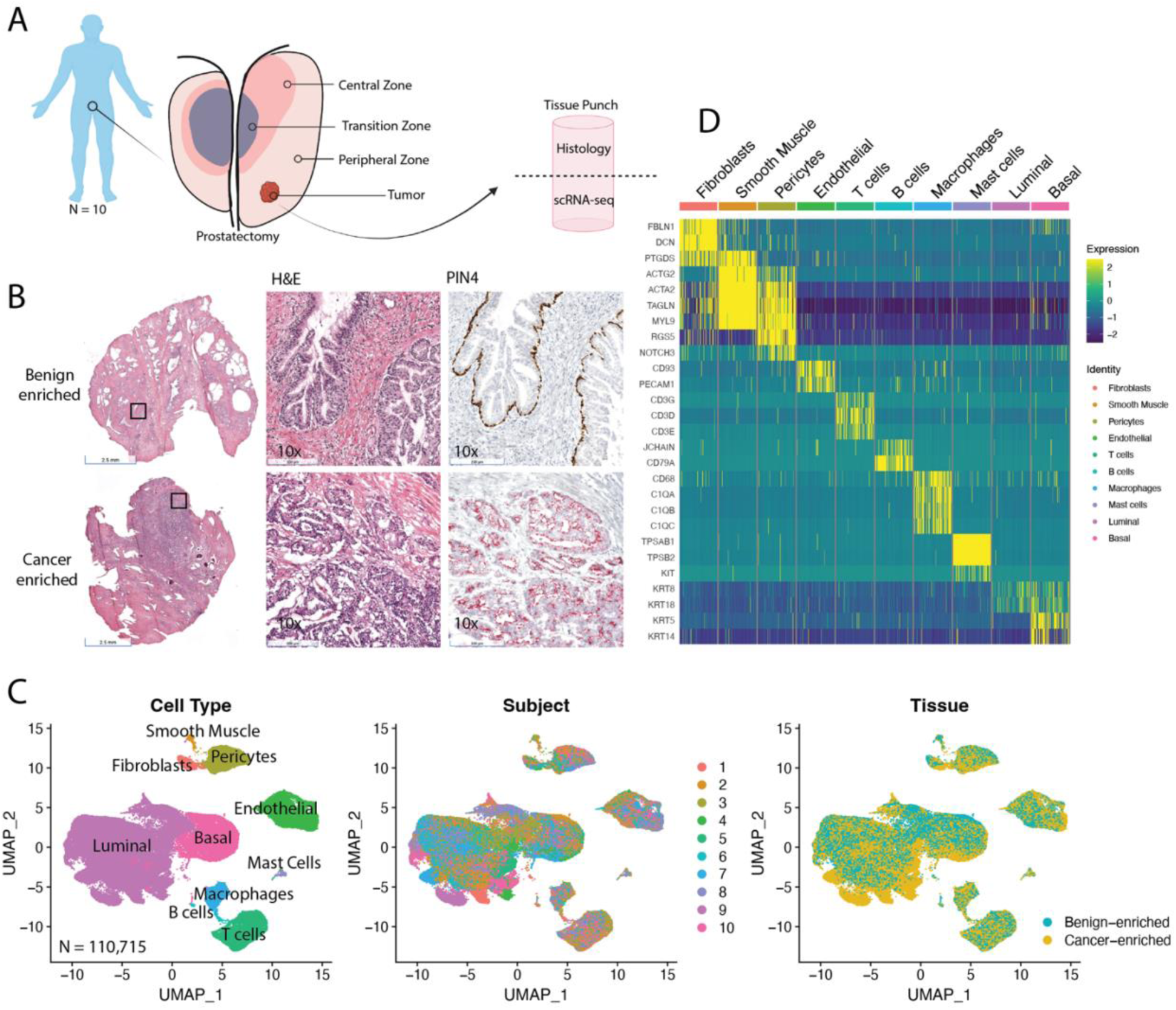
Single-cell RNA sequencing (scRNA-seq) of prostatectomy tissues collected from patients diagnosed with primary prostate cancer. A) For each prostatectomy (N = 10), tissue punches were collected from each zone of the prostate as well as the tumor site (4 punches per subject). For each tissue punch, part of the tissue was processed for scRNA-seq from freshly dissociated cells, and the remaining half was frozen and sectioned for histology, IHC, and in situ hybridization. B) Representative examples of hematoxylin and eosin (H&E) staining of fresh frozen peripheral zone tissue punches that were benign enriched or cancer enriched. The far right panels are immunohistochemical PIN4 staining. Brown marks tumor protein P63 (TP63) and high molecular weight cytokeratin expressed in basal epithelial cells. Red marks alpha-methylacyl-CoA racemase (AMACR), which is expressed in carcinoma and prostatic intraepithelial neoplasia (PIN). C) Dimensionality reduction (uniform manifold approximation and projection, UMAP) and clustering analysis of scRNA-seq libraries from peripheral zone (PZ) tissue showed cells clustering by recognized cell types. Subject and cancer-enriched zones of luminal cells suggest inter-individual and intra-tumor heterogeneity in cancer cells. D) Heatmap of cell type marker genes show cluster-specific expression.

### Defining benign and neoplastic epithelial cell states and associated gene expression and copy number alterations

We next defined the epithelial cell states in the prostate tissues with greater granularity by subsetting the luminal and basal clusters and repeating dimensionality reduction and clustering analyses. We used a combination of differential gene expression analysis, well-known gene expression markers of known epithelial cell types, and inferred copy number alterations derived from the gene expression data to further refine the identification of neoplastic cells harboring copy number alterations known to be recurrent in human prostate cancer (see Supplementary Information for details) ^18, 33–37^. With these analyses, we identified benign basal and luminal cells, proliferative atrophy/intermediate cells, which are similar to what others have previously referred to as club/intermediate cells ^38–41^, and eight distinct PIN/cancer clusters that we referred to as Cancer-1 through Cancer-8 (Figures 2A-B, S1). These cancer clusters showed strong enrichment of specific subjects, again consistent with the known inter-individual molecular heterogeneity of prostate cancer. In contrast, the benign basal, luminal and proliferative atrophy/intermediate clusters had representation from all ten subjects, with little inter-individual heterogeneity (Figures 2A-B).

**Figure 2.**
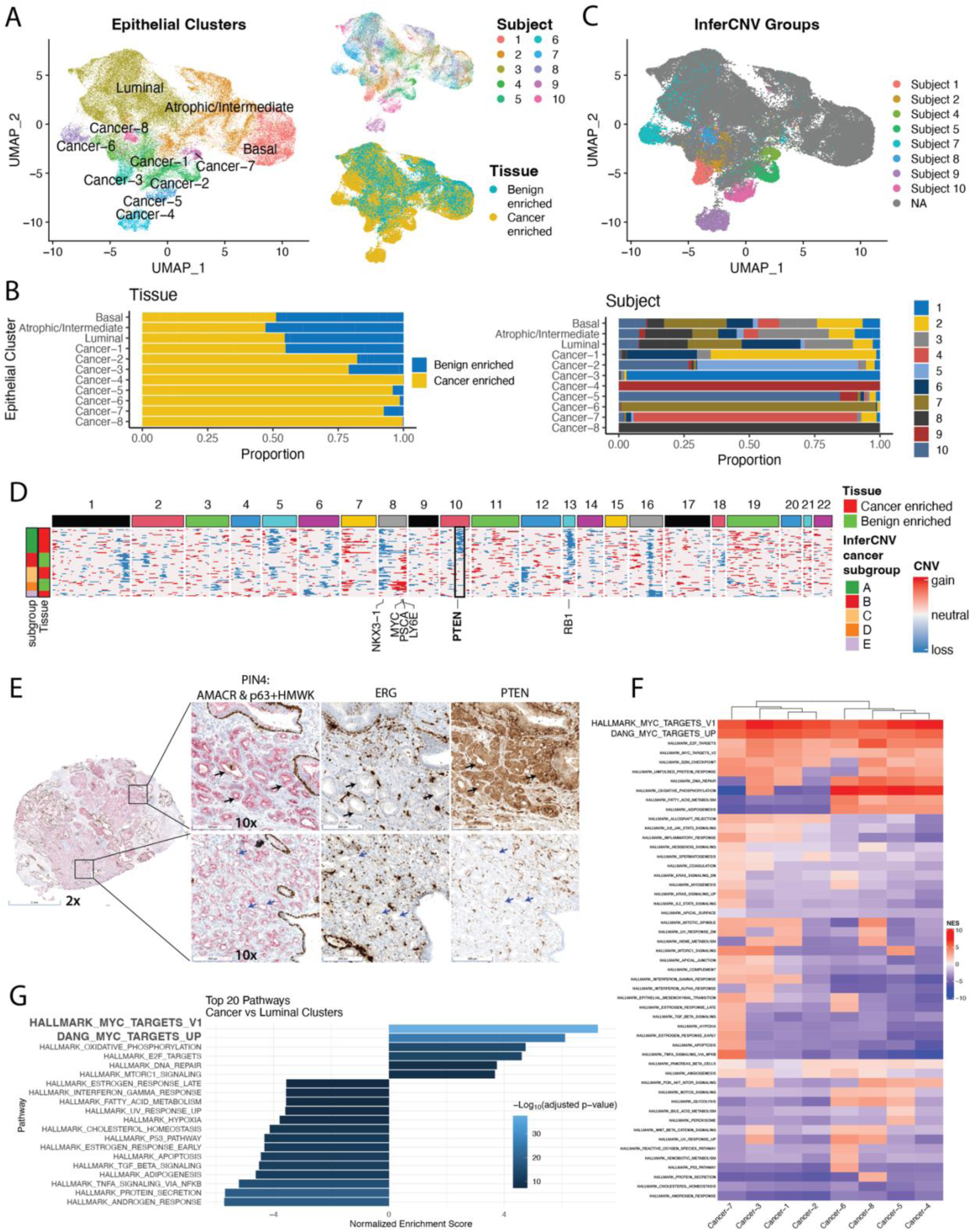
Prostate cancer inter-individual and intra-tumor heterogeneity. A) Epithelial clusters were subsetted, and dimensionality reduction (UMAP) and clustering analysis were repeated. Benign and cancer cells clustered separately. B) Stacked bar plots showing the proportion of cells for each cluster that are from (left panel) benign or cancer-enriched libraries and (right panel) which subject (1-10). C) UMAP indicating cells from inferred copy number variation (InferCNV) analysis harboring cancer-associated mutations by subject. D) Heatmap from inferCNV analysis of cancer cells from Subject 2 indicating regions with inferred CNV gain (red) and loss (blue). Cancer subgroup A shows 10q loss, while the remaining subgroups show 10q is intact, suggesting heterogeneous loss of PTEN (black box). E) Example of intra-tumor heterogeneity from Subject 2. Immunohistochemical for frozen sections for PIN4, ERG, and PTEN staining shows cancer cells in top panels are negative for ERG and positive for PTEN. In contrast, lower panels show another group of cancer cells that are ERG-positive but PTEN-negative. F) Heatmap of normalized enrichment scores (NES) from gene set enrichment analysis (GSEA) of Hallmark collection and “Dang MYC targets up gene set” comparing each cancer cluster with the luminal cluster. G) Plot showing NES of top 20 pathways by adjusted p-value, comparing aggregated cancer clusters with the luminal cluster by GSEA.

The PIN/cancer clusters showed distinct gene expression programs and copy number alterations consistent with known prostate cancer subtypes that were enriched in specific subjects (Figures 2A-B, S1, Supplementary Information), including ERG fusion-positive cancers (Cancer-2, Cancer-4, and Cancer-5 clusters; enriched in subjects 5, 9, and 10 respectively) and SPINK1 positive cancers (Cancer-1, Cancer-3, Cancer-6, Cancer-7, and Cancer-8; enriched in subjects 2, 1, 7, 4, and 8 respectively) (Figure S1B) ^15–17, 42, 43^. Consistent with known patterns in human prostate cancer, we observed ERG-positive cases with retained PTEN expression, ERG-positive cases with complete PTEN loss, and ERG-negative cases with PTEN-intact expression (Figure S2) ^16, 17, 44, 45^. Furthermore, inferred copy number alteration patterns showed significant heterogeneity across subjects, consistent with known regions of recurrent copy number alteration in primary prostate cancer, and strong parallels with the gene expression derived clustering (Figure 2C, Supplementary Figures S3-5, Supplementary Information) ^17, 46^.

In addition to the striking inter-individual heterogeneity in the cancer clusters, the scRNA-seq-derived copy number analysis could also capture intra-individual tumor heterogeneity. For example, in subject 2, a subset of luminal cells had 10q loss, a common CNV in prostate cancer consistent with PTEN loss ^47–50^, while other cancer cells from this subject did not harbor 10q/PTEN loss (Figures 2D and S3B). Tissue staining from subject 2 showed heterogeneous loss of PTEN, with one group of cancer cells with PTEN expression intact and another group with loss of PTEN (Figure 2E), consistent with the intratumoral heterogeneity of PTEN deletion identified by the inferCNV analysis.

These in situ tissue analyses corroborated that our scRNA-seq analyses could capture the well-known inter- and intra-individual tumor heterogeneity in prostate cancer. Additionally, our scRNA-seq analyses showed that there was very little inter-individual heterogeneity in benign basal and luminal epithelial cells, proliferative atrophy/intermediate cells, and stromal cells.

### MYC activation is a common denominator across the known molecular heterogeneity of prostate cancer

To identify pathways dysregulated in cancer clusters 1-8, we performed gene set enrichment analysis (GSEA) comparing benign luminal epithelial cells with each of the eight cancer clusters using the Hallmark gene set collection. As expected, there was variation in the upregulation and downregulation of several gene sets (Figure 2F), consistent with the heterogeneity we observed in our multimodal investigation of prostate cancer tissues by scRNA-seq clustering analysis, inferred CNVs, and *in situ* tissue staining. Two gene sets were universally downregulated, including androgen response and cholesterol homeostasis. Three Hallmark gene sets were upregulated in all eight cancer clusters: MYC Targets V1, MYC Targets V2, and E2F Targets. Aggregating cancer clusters 1-8 and performing differential expression and pathway analysis of this aggregated cancer cluster against benign luminal cell clusters revealed that the Hallmark MYC Targets V1 gene set was the top upregulated pathway and androgen response the top downregulated pathway in the cancer clusters (Figure 2G, and Table S4). The universal upregulation of MYC targets and downregulation of the androgen response pathway across all cancer clusters are consistent with recent reports characterizing the antagonistic relationship between MYC and androgen response genes ^51–53^. Including the Dang MYC Targets Up gene set in GSEA ^54^ reaffirmed significant upregulation of MYC targets across all cancer clusters (Figures 2F-G). To confirm that upregulated MYC activity is prevalent across prostate cancer subtypes, we examined the TCGA prostate cancer dataset, which was instrumental in defining molecular subtypes of prostate cancer ^16^. We found upregulated MYC activity in the vast majority of prostate cancer samples, regardless of Gleason score and molecular subtype (Figure S6).

Additionally, we observed elevated expression of MYC Targets V1 genes in the basal, atrophic/intermediate, and cancer clusters compared to benign luminal epithelial clusters (Figures S7A-F). Validation studies with *in situ* tissue staining confirmed the over-expression of nuclear MYC in prostate cancer cells and basal cells from benign glands compared to normal luminal epithelial cells (Figure S7G). These findings are consistent with prior immunostaining studies showing that *MYC* expression is common in prostatic basal cells, negative in normal luminal cells, somewhat upregulated in atrophic luminal/intermediate cells, and its overexpression is a highly frequent and early event in prostate cancer ^22^.

### MYC activity reprograms neighboring benign epithelial cells in the TME

It is difficult to assess the causal sequelae of MYC activation in human tissues. Therefore, to investigate the downstream consequences of MYC activation in the prostate TME, we modeled MYC activation in mouse prostates using transgenic Hi-Myc mice ^51, 55–58^. Because the 6-month timepoint marks a critical switch from precursor to invasive carcinoma in the Hi-Myc model, we rigorously assessed the single-cell transcriptomes in each prostate lobe (anterior, dorsal, lateral, and ventral) of the MYC-driven mouse model of prostate cancer (Hi-Myc) in both FVB/NJ and C57BL/6J mouse strains aged 6 months (Figures 3A-C, Supplemental Figure S8, Supplemental Information). As a reference, age-matched wild-type (WT) animals from our previously published single-cell atlas of the mouse prostate by lobe and strain were included in the analysis ^59^.

**Figure 3.**
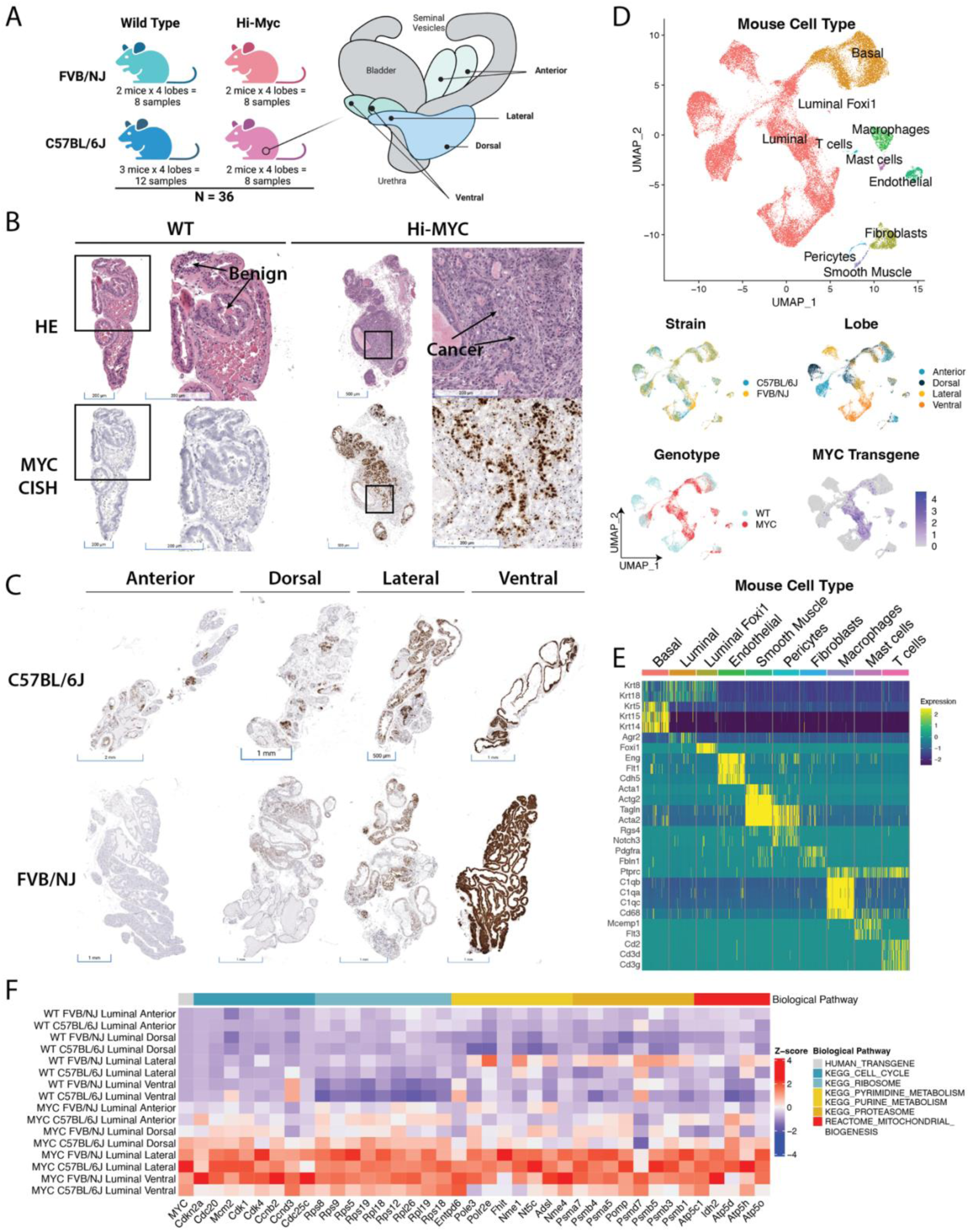
Single-cell RNA-seq and in situ analysis of Hi-Myc mouse model of prostate cancer. A) Tissue was collected from each prostate lobe (anterior, dorsal, lateral, and ventral) of the MYC-driven mouse model of cancer (Hi-Myc) in both FVB/NJ (N = 2) and C57BL/6J (N =2) mouse strains, as well as age-matched wild type animals (FVB/NJ N =2, C57BL/6J N = 3). B) H&E (upper panels) and chromogenic in situ hybridization (CISH, lower panels) staining of human MYC transgene in FFPE mouse prostate tissues from 6-month-old Hi-Myc and age-matched wild-type (WT) mice. C) Representative CISH staining of human MYC transgene in each lobe of Hi-Myc mouse prostate from C57BL/6J and FVB/NJ mice. D) UMAP of mouse scRNA-seq by cell type, strain, lobe, genotype, and *MYC* transgene expression. E) Heatmap of each cluster and cell type marker gene expression. F) Heatmap showing expression of genes positively correlated with *MYC* transgene expression in luminal cell populations stratified by genotype, strain, and lobe. Genes are grouped by associated biological pathways. The first column shows the scaled expression of the *MYC* transgene.

Dimensionality reduction and clustering analysis showed that cells grouped primarily by known cell types (Figures 3D-E, S9A). Consistent with previously reported single-cell atlases of the mouse prostate ^59, 60^, luminal epithelial cells from WT mice subclustered by lobe and strain (Figure S9A). However, Hi-Myc luminal cells clustered by genotype and lost strain and lobe-specific clustering, with the exception of the luminal cells from the ventral lobe (Figures 3D, S9). Genes positively correlated with *MYC* transgene expression in luminal cells were associated with several biological pathways linked to MYC activation (Figure 3F), including cell cycle progression, ribosome biogenesis, RNA processing, protein degradation, and mitochondrial function ^54, 61–63^.

To investigate the MYC-driven changes in the epithelial population, we subsetted our analysis to the luminal and basal epithelial cells and identified four epithelial clusters disproportionately enriched in Hi-Myc mice (Figures 4A-C, Table S5, Supplemental Information). Two of the epithelial clusters that were strongly enriched in the Hi-Myc compared to WT animals showed strong expression of the human MYC transgene and upregulation of MYC targets (Figure 4D). Given the in situ analysis of MYC transgene expression coinciding exclusively in PIN and cancer lesions in Hi-Myc mice (Figures 3B, C), the cells of these two clusters represent the precursor and invasive carcinoma cells in the Hi-Myc animals. We therefore termed these clusters Luminal MYC 1 and Luminal MYC 2 and noted that they were enriched in the dorsal/lateral vs. ventral lobes, respectively (Figure 4C).

**Figure 4.**
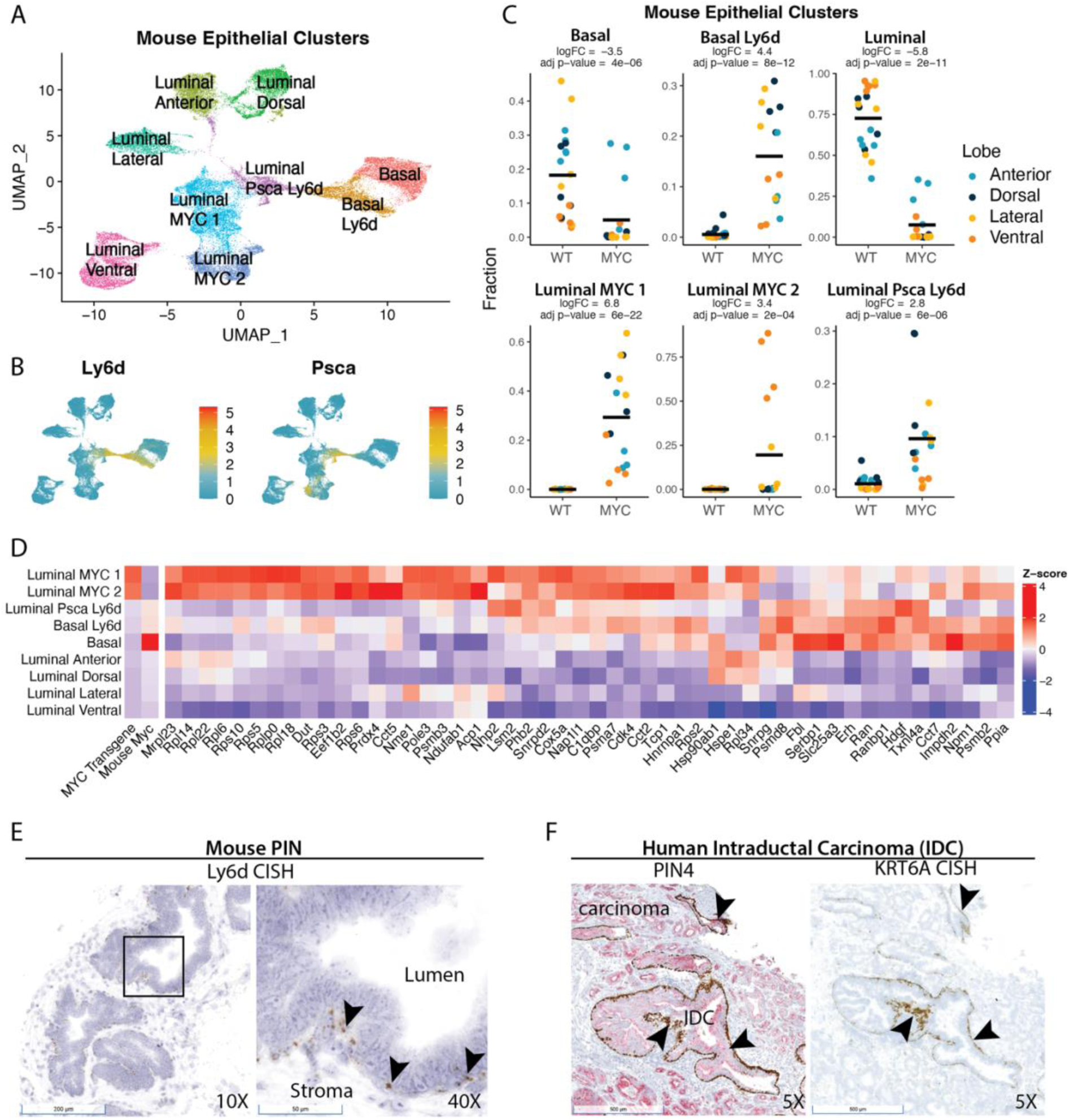
Cascading changes in the epithelial population in the tissue microenvironment (TME) of *MYC*-driven prostate cancer. A) UMAP of mouse prostate scRNA-seq subsetted to epithelial clusters. B) UMAPs of *Defb29*, *Timp1*, *Ly6d,* and *Psca* show cluster-enriched gene expression. C) Scatter plots showing the proportion of epithelial clusters for each sample. Each dot represents a sample colored by lobe (anterior, dorsal, lateral, ventral). Bar represents the mean expression of samples for each genotype. Statistics were generated by comparing WT and Hi-Myc samples using regression analysis in scRNA-seq with multiple samples (RAISIN) cell proportions test. D) Heatmap showing expression of endogenous *Myc*, transgene *MYC*, and Hallmark MYC targets V1 leading edge genes from GSEA for each epithelial cluster. E) Representative example of *Ly6d* CISH staining basal compartment in PIN gland of Hi-Myc FFPE prostate tissue. F) Example of *KRT6A* CISH staining basal compartment of intraductal carcinoma in human prostate cancer from frozen tissue sections.

Surprisingly, the other two clusters, *Ly6d*-expressing luminal and basal cells, did not show robust expression of the human MYC transgene despite being significantly enriched in the Hi-Myc mice. This suggested that MYC activation in neoplastic luminal cells led to the induction of these *Ly6d*-expressing basal and luminal cells in the surrounding microenvironment. We therefore further explored these cell clusters (Supplemental Information) and sought to identify parallels in the human prostate cancer tissues.

Gene expression analysis showed that *Krt6a* was one of the genes uniquely upregulated in Ly6d-expressing luminal and basal cells and strongly co-expressed with *Ly6d* (Supplemental Figures S10A-B). Expression analysis of epithelial cells from prostatectomy samples revealed that a subset of basal cells expressed KRT6A and LY6D, suggesting the mouse Basal Ly6d cells enriched in Hi-Myc mouse prostates were also observed in the human prostate (Figure S10C). Chromogenic in situ hybridization (CISH) for *Ly6d* in Hi-Myc mouse prostate tissues revealed a subset of PIN lesions expressing Ly6d in the basal compartment (Figure 4E). Likewise, CISH staining in human prostatectomy tissues revealed that *KRT6A* was expressed primarily in a subset of basal cells of prostate glands, including PIA, PIN, intraductal carcinoma, and to a much lesser extent, a subset of intermediate/atrophic cells in PIA (Figure 4F, Supplemental Figure S11).

Although the Ly6d basal cluster was highly enriched in the Hi-Myc prostate, the lack of *MYC* transgene expression and in situ tissue staining pattern of *Ly6d* in mouse prostates and *KRT6A* in human prostates suggest that this cell type may be induced by MYC-transformed neoplastic cells through paracrine signaling (Figures 4C-D). Inferred ligand-receptor paired interactions revealed that the Luminal MYC 1 cluster had the strongest inferred interactions with the Basal Ly6d cluster, expressing several of the top ligands targeting the Basal Ly6d cluster, including the type I interferon ligand, interferon kappa (*Ifnk*) (Figures 5A-B) ^64, 65^. Interestingly, expression of the type I interferon receptor subunits *Ifnar1* and *Ifnar2* in the Basal Ly6d cells was positively correlated with the activity of the interferon regulatory factor IRF5 as measured by expression of its downstream targets (Figures 5C-D). In line with upregulated IRF5 activity, the Basal Ly6d cluster also had increased *Irf5* expression (Supplemental Figure S12). Consistent with activated interferon signaling in the TME, the Luminal MYC 1 and Basal Ly6d clusters were both found to be upregulated in interferon alpha and gamma response pathways (Figure 5E). These cell communication network and pathway analyses suggested that MYC-expressing neoplastic cells induce enrichment of neighboring Ly6d-expressing epithelial cells in part via type I interferon signaling and other ligand-receptor interactions (Figures 5A-E, Supplementary Information).

**Figure 5.**
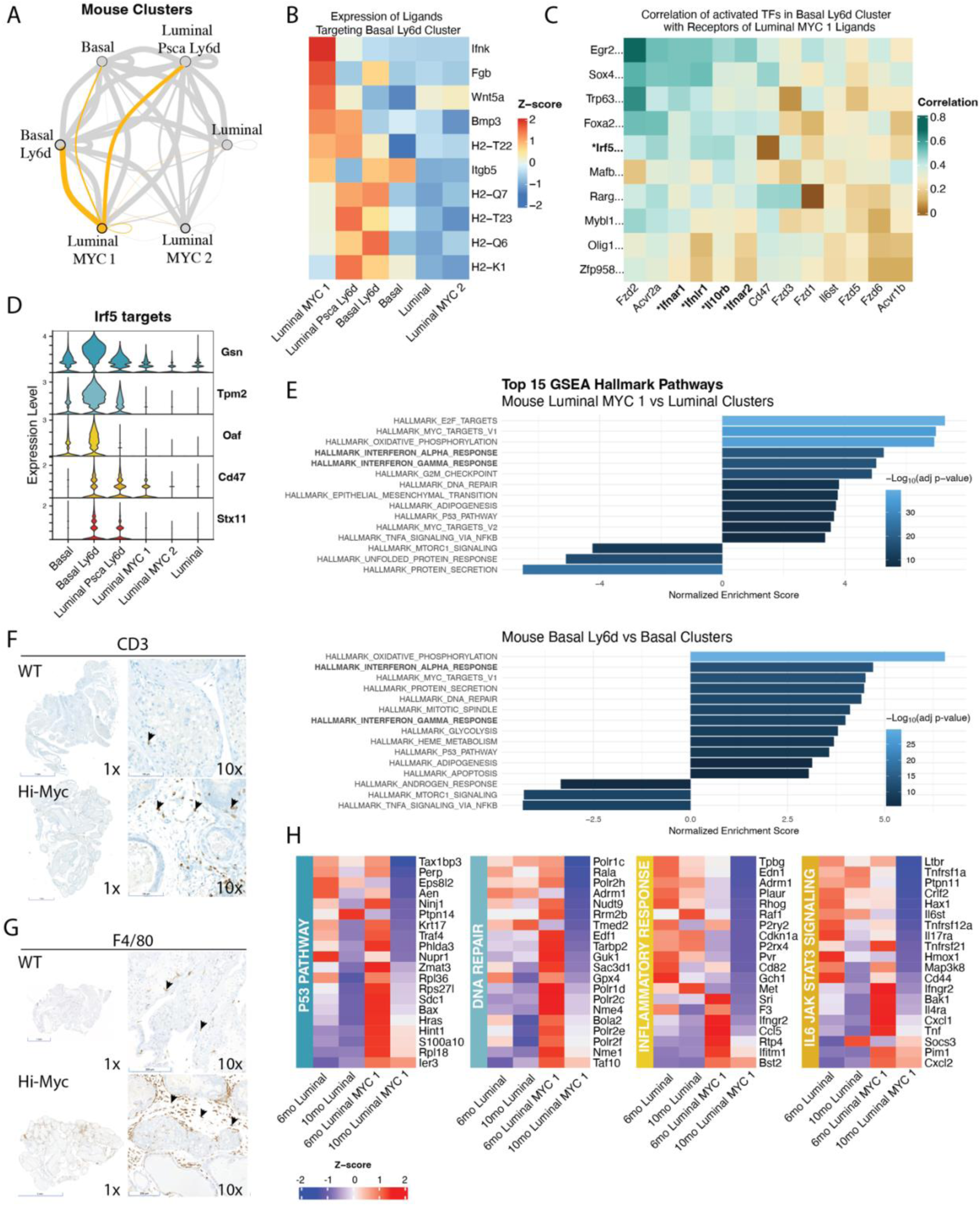
Signaling among epithelial clusters in TME of *MYC*-driven prostate cancer. A) Cell communication plot of epithelial clusters from inferred ligand-receptor-transcription factor network analysis (Domino). Nodes represent individual clusters and are scaled based on ligand expression. Edges indicate signaling between two clusters and are weighted based on the strength of signaling. The edge color matches the cluster expressing the ligand. The Luminal Timp1 cluster is marked in yellow, while all other clusters are colored grey to highlight specific interactions from the Luminal Timp1 cluster to all other epithelial clusters. B) Heatmap showing which epithelial clusters express the top ligands targeting receptors expressed on the Basal Ly6d cluster from inferred ligand-receptor pair interactions. C) Heatmap showing the correlation of transcription factor activity scores (from SCENIC analysis) and receptor expression. Transcription factors were selected based on top activated scores in the Basal Ly6d cluster. The receptors represented in this heatmap are known to bind to the top ligands targeting Basal Ly6d. D) Expression of downstream targets of the transcription factor interferon response factor 5 (IRF5) in epithelial clusters. E) Plot showing NES of top 15 pathways by adjusted p-value, comparing Luminal and Luminal MYC 1 clusters (top panel) and Basal and Basal Ly6d clusters (bottom panel). Immunohistochemical staining in WT and Hi-Myc mouse prostate tissues at 6 for F) CD3 to mark T cells and G) F4/80 to mark macrophages. H) Heatmaps of top 20 genes upregulated in the Luminal Timp1 cluster at 6 months and downregulated at 10 months. Genes are grouped by associated gene sets from the GSEA Hallmark collection.

### Proinflammatory to immunosuppressive switch in MYC-expressing luminal cluster accompanying progression to invasive cancer

In addition to upregulated interferon response pathways, GSEA comparing Luminal and Luminal MYC 1 cells also revealed upregulation of another pro-inflammatory pathway, tumor necrosis factor alpha (TNFA) signaling via nuclear factor kappa-light-chain-enhancer of activated B cells (NF-κB) (Figure 5E). In both FVB/NJ and C57BL/6J strains, we observed a significant enrichment of various immune cells, including mast cells, T cells, and *Trem2*-expressing macrophages in Hi-Myc prostates compared to age-matched WT prostates (Supplemental Figure S13, Supplemental Information). Prior studies have suggested TREM2 Macrophages have an immunosuppressive role in the TME ^66, 67^. Immunostaining for T cells (CD3) and macrophages (F480) in WT and Hi-Myc mouse prostates showed enrichment of T cells and macrophages in the prostate TME of Hi-Myc mice (Figures 5F-G), consistent with the upregulation of inflammatory pathways observed by GSEA in luminal epithelial cells (Figure 5E).

To better understand the dynamics of the epithelial and immune cell phenotypes in response to MYC activation and cancer progression over time, we expanded our analysis to include a later time point (10 months +/-6 weeks) of dorsal and lateral lobes from Hi-Myc and aged-matched WT prostates from FVB/NJ mice. The FVB/NJ dorsal and lateral lobes in Hi-Myc mice are are known to progress to more penetrant invasive cancer phenotypes by the 10 month time point and would therefore allow us to understand the dynamics of the TME during cancer progression (Figure S8). We merged scRNA-seq datasets from the dorsal and lateral lobes of Hi-Myc and aged-matched WT prostates from FVB/NJ mice aged 6 months and 10 months and integrated by age (Figures S14A-B, Table S6). GSEA of MYC-expressing Luminal MYC 1 cluster comparing 10-month and 6-month mice revealed a shift in multiple pathways (Figure S14C). In particular, the Luminal MYC 1 cluster in 10-month old mice downregulated several genes associated with critical pathways that serve as gatekeepers of malignant transformation, including P53 and DNA repair (Figures 5H, S14C). Strikingly, the Luminal MYC 1 cluster was also downregulated for several inflammation-related genes, including inflammatory response and IL6 JAK STAT3 signaling (Figures 5H, S13C). Collectively, these pathway analyses suggest that a cancer-cell intrinsic pro-inflammatory to immunosuppressive switch coincides with pathway alterations associated with cancer progression.

We next examined whether the cancer cell-intrinsic switch from pro-inflammatory to immunosuppressive programs was accompanied by alterations in the composition of the immune microenvironment between the 6-month, precursor-enriched, and 10-month, invasive cancer-enriched time points. We first defined the major immune cell clusters (Figures 6A-B) and noted several distinct differences between the two time points. First, we observed a distinct enrichment in the 10-month vs. 6-month timepoint of multiple cell populations known to have immunosuppressive function, including TREM2 positive macrophages, myeloid-derived suppressor cells (MDSCs), and regulatory T cells (T regs), which could also be confirmed by in situ hybridization analysis of the mouse prostate tissues (Figure 6C, Supplemental Figure S15). In the human prostatectomy dataset, we observed a subset of T cells that express regulatory T-cell-associated markers in a gene signature analysis (UCell) (Figures 6D-F). We also observed that a subset of macrophages expressed TREM2 (Figure 6G), suggesting similar immune composition changes driven by MYC oncogene activity in mice are observed in human prostate cancer. Notably, a recent scRNA-seq prostate cancer study focused on invasive cribriform carcinoma and intraductal carcinoma also reported increased TREM2 macrophages in the TME compared to benign tissues ^68^.

**Figure 6.**
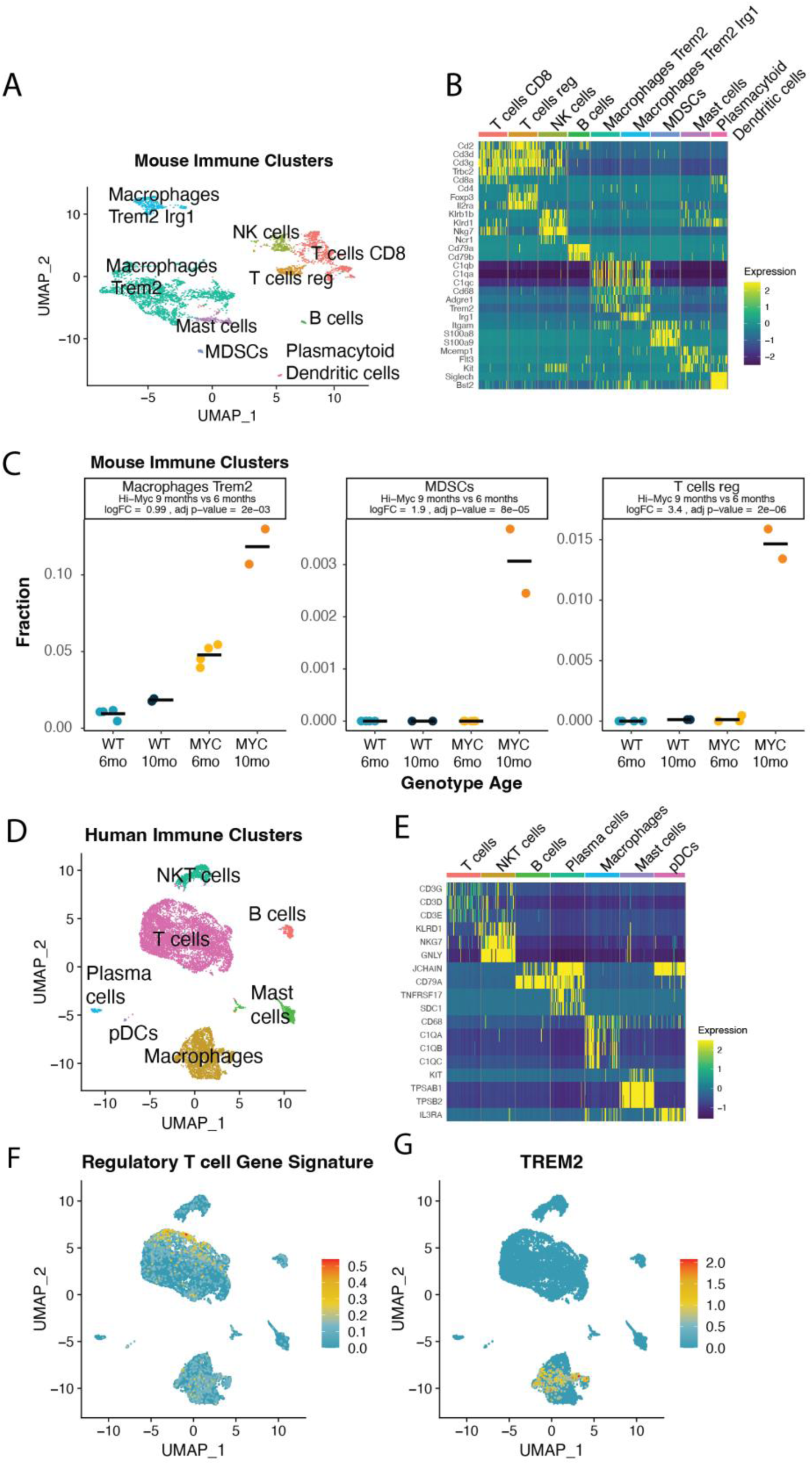
Immune cell populations enriched in the TME of *MYC*-driven prostate cancer. A) Heatmap of immune cell type marker genes show cluster-specific expression in merged scRNA-seq data from dorsal and lateral lobes of FVB/NJ mice from WT and Hi-Myc at 6 months and 10 months. B) Immune cell clusters were subsetted, and dimensionality reduction (UMAP) and clustering analysis were repeated. C) Scatter plots showing the proportion of select immune cell populations for each sample. Each dot represents a sample colored by genotype and age. Bar represents the mean expression of samples for each group. Statistics were generated by comparing Hi-Myc samples at 6 and 10 months using RAISIN’s cell proportions test. D) UMAP of human immune cell clusters from prostatectomy scRNA-seq dataset. E) Heatmap of immune cell type marker gene expression of immune cells from prostatectomy samples. Expression of F) regulatory T cell gene signature (*IL2RA*, *FOXP3*, *CD4*, *IKZF2*, *CCR4*, *CTLA4*) and G) *TREM2* in human immune cell clusters.

### Stromal cells are significantly altered in the TME of MYC-driven prostate cancer

In the merged 6-month and 10-month FVB/NJ dorsal and lateral datasets, the stromal populations included smooth muscle, pericytes, endothelial, glial, and fibroblast cells (Figure 7A, Supplemental Figures S16A-B, Table S7). Hi-Myc animals showed increased fractions of endothelial cells, consistent with induction of neovasculature in MYC-driven neoplasia (Figure 7B).

**Figure 7.**
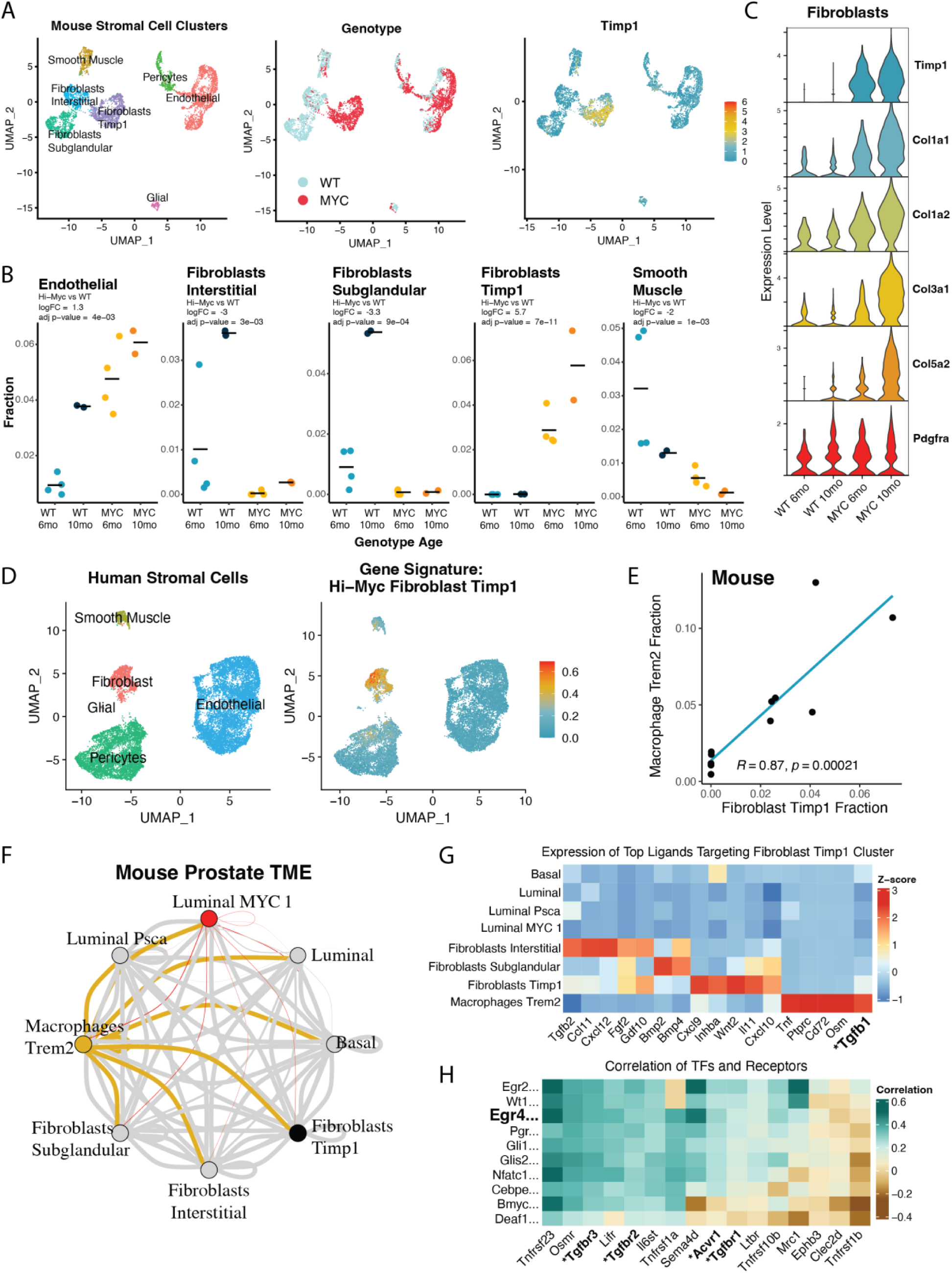
Stromal reprogramming in the TME of *MYC*-driven prostate cancer. A) Immune cell clusters were subsetted from merged scRNA-seq data of dorsal and lateral lobes of FVB/NJ mice from WT and Hi-Myc at 6 months and 10 months, and dimensionality reduction (UMAP) and clustering analysis were repeated. A fibroblast subcluster enriched in Hi-Myc tissue specifically expresses *Timp1*. B) Violin plot showing expression of *Pdgfra*, a marker of fibroblasts, *Timp1*, and fibrosis-associated collagens (*Col1a1*, *Col1a2*, *Col3a1*, *Col5a2*) by genotype (WT and Hi-Myc) and age (6 months and 10 months) in fibroblast cells. C) Scatter plots showing the proportion of select stromal cell populations significantly altered in Hi-Myc compared to WT. Each dot represents a sample colored by genotype and age. Bars represent the mean expression of samples for each group. Statistics were generated by comparing WT and Hi-Myc samples using RAISIN’s cell proportions test. D) UMAP of human stromal cell clusters from prostatectomy samples. The right panel shows the expression of the Hi-Myc Fibroblast Timp1 gene signature (*TIMP1, MFAP5, SERPINA3, IGF1, SFRP1, MMP2, SERPINF1, COL1A1, COL5A2, COL3A1*) in human stromal clusters. E) Scatter plot showing proportion of Macrophages Trem2 and Fibroblast Timp1 clusters for each sample with corresponding Pearson correlation coefficient and p-value. F) Cell communication plot of inferred ligand-receptor-transcription factor network analysis (Domino) of epithelial, macrophage, and fibroblast clusters from FVB/NJ aggregated scRNA-seq data. The Luminal Timp1 cluster is marked in red, the Macrophages Trem2 cluster is marked in yellow, the Fibroblast Timp1 node is marked in black, and all other clusters are colored grey. G) Heatmap showing which clusters express the top ligands targeting the Fibroblast Timp1 cluster from inferred ligand-receptor pair interactions. The ligand transforming growth factor beta 1 (*Tgfb1*) is bolded. H) Heatmap showing the correlation of transcription factor activity scores (derived from SCENIC) and receptor expression. Transcription factors were selected based on top activated scores in the Fibroblast Timp1 cluster. Receptors for TNF, PTPRC, CD72, OSM, and TGFB1 are shown. Transcription factor early growth response 4 (EGR4) and genes encoding the receptors of TGFB1 (*Tgfbr1, Tgfbr2, Tgfbr3,* and *Acvr1)* are bolded.

The fibroblasts sub-clustered into three groups, including two fibroblasts previously characterized in WT mouse prostates, *Sult1e1*-expressing interstitial fibroblasts and *Rorb*-expressing subglandular fibroblasts ^59, 69^, and a novel *Timp1*-expressing fibroblast cluster (Fibroblast Timp1) that was highly enriched in Hi-Myc mice. This cluster showed significantly increased expression of fibrosis-associated collagens *Col1a1*, *Col1a2, Col3a1, and Col5a2* ^70–72^, consistent with a reactive stroma and the desmoplastic pan cancer-associated fibroblast gene expression signature (pan-dCAF) ^30, 73^ (Figures 7C, Supplemental Figure S17, Supplemental Information). More broadly, the enrichment of fibrosis-associated extracellular matrix (ECM) gene expression in the TME of prostate cancer is consistent with a desmoplastic response, a feature associated with invasive carcinoma in tissues ^74–76^. Genes differentially expressed in the Fibroblast Timp1 cluster in mice were evaluated in the human stroma. Gene signature analysis revealed that a subset of fibroblasts in human prostatectomy samples was enriched for the Fibroblast Timp1 gene signature (Figures 7D, S16C-D, Table S9).

In the mouse prostate TME, this increase in the Fibroblast Timp1 CAFs was accompanied by decreases in the normal interstitial and subglandular fibroblasts and smooth muscle clusters in the Hi-Myc mice (Figure 7B). Similar stromal changes were also observed in C57BL6/J Hi-Myc mice, with enrichment of endothelial cells and Timp1-expressing fibroblasts and a reduced proportion of interstitial fibroblasts (Supplemental Figure S18, Table S8). Intriguingly, these Fibroblast Timp1 CAFs increased in the 10-month invasive cancer enriched compared to 6-month precursor-enriched tissues (Figure 7B).

Moreover, the fraction of cells in the Fibroblast Timp1 cluster was significantly correlated with the fraction of cells in the Macrophage Trem2 cluster (Figure 7E), suggesting that these two clusters may be connected through paracrine signaling. Inferred ligand-receptor-transcription factor network analysis in the mouse dataset showed strong inferred interactions of the Macrophage Trem2 cluster with the Fibroblast Timp1 cluster (Figure 7F). In our scRNA-seq data, many ligands targeting the Fibroblast Timp1 cluster were expressed in the Macrophage Trem2 cluster, including transforming growth factor beta 1 (*Tgfb1*) (Figure 7G). Notably, early growth response 4 (EGR4) activation scores positively correlated with TGFB1 receptors *Tgfbr1*, *Tgfbr2*, *Tgfbr3*, and activin A receptor type 1 (*Acvr1*) in the Fibroblast Timp1 cluster (Figure 7H). Downstream targets of EGR4 include *Timp1* and the fibrosis-associated collagens *Col1a1*, *Col1a2*, *Col3a1*, and *Col5a2*. Importantly, TGFB signaling is a key instigator of tissue fibrosis and is thought to be mediated by immune cells ^77, 78^. Indeed, scRNA-seq analysis coupled with in situ tissue validation is consistent with a sequence of events involving infiltration and expansion of *Tgfb1* expressing macrophages in tissues with MYC oncogene expressing luminal cells, coinciding with dramatic stromal remodeling enriched for a unique fibroblast population with robust collagen expression.

## DISCUSSION

Primary prostate cancer is a heterogeneous disease with multiple genomic alterations driving carcinogenesis. For example, common alterations include fusion and overexpression of E26 transformation-specific (ETS) transcription factors, as well as mutations in speckle type BTB/POZ protein (*SPOP*), forkhead box A1 (*FOXA1*), isocitrate dehydrogenase (NADP(+)) 1 (*IDH1*), and phosphatase and tensin homolog (*PTEN*) ^1, 15, 16, 79, 80^. Yet, despite the underlying genomic heterogeneity typical of this disease, there are several key characteristics of the TME shared across most primary prostate cancer. The glandular architecture characteristic of the normal prostate, which consists of cuboidal basal cells aligning the perimeter of differentiated columnar epithelial cells, devolves in carcinoma, with cancer cells invading the surrounding fibromuscular stroma ^81–83^. In digital rectal exams, areas of the prostate suspected of harboring cancer cells can stiffen and feel indurated ^84^, likely due to the desmoplastic response driven by an inflammatory reactive stroma characterized by infiltrating immune cells and increased deposition of extracellular matrix proteins ^30^.

Our single-cell and complementary in situ tissue analysis of prostatectomy tissues collected from 10 patients diagnosed with primary prostate cancer are consistent with the notion that prostate cancer is a heterogeneous disease. Yet, despite this tumor heterogeneity, we observed MYC activation as a common denominator across these cancer clusters. This observation was supported when performing differential gene expression and pathway analysis of the publicly available TCGA dataset of primary prostate cancer ^16^.

Evaluating the direct effects of MYC activation in the prostate TME is challenging in human tissues. Therefore, we modeled MYC activation using the transgenic Hi-Myc mouse model of prostate cancer ^55, 56^ and used computational approaches to evaluate changes in the TME. To account for potential strain-specific effects, we evaluated the Hi-Myc mouse model of prostate cancer in both FVB/NJ and C57BL/6J mouse strains. Key insights from mouse model studies were evaluated in prostatectomy tissue samples to confirm that observations from mouse studies were not a mouse model-specific phenomenon but also relevant to human prostate cancer.

Based on these analyses, we found that MYC activation in neoplastic cells sets off a cascade of alterations that fundamentally alters the composition and transcriptional programs of surrounding cell types in the TME (Figure 8). Initially, MYC activation is associated with an induction of a pro-immunogenic transcriptional program in a precursor-enriched state. This leads to propagation of interferon and inflammatory mediators that induce a new basal cell state, likely through engagement of type I interferon receptors. These newly programmed basal Ly6d/Krt6a-expressing cells in turn propagate a pro-immunogenic program with resulting influx of pro-inflammatory cells including macrophages, mast cells, and T cells. Over time, the MYC-activated luminal neoplastic cells in the precursor state undergo a switch to significantly downregulate the pro-immunogenic transcriptional programs alongside downregulation of other key tumor suppressor pathways during progression to invasive cancer. This change is accompanied by an enrichment of immunosuppressive cells in the microenvironment including MDSCs, Tregs, and TREM2+ macrophages. This immunosuppressive microenvironment is consistent with the well-established observation that the prostate cancer TME is immunologically cold ^26–28, 85^.

**Figure 8.**
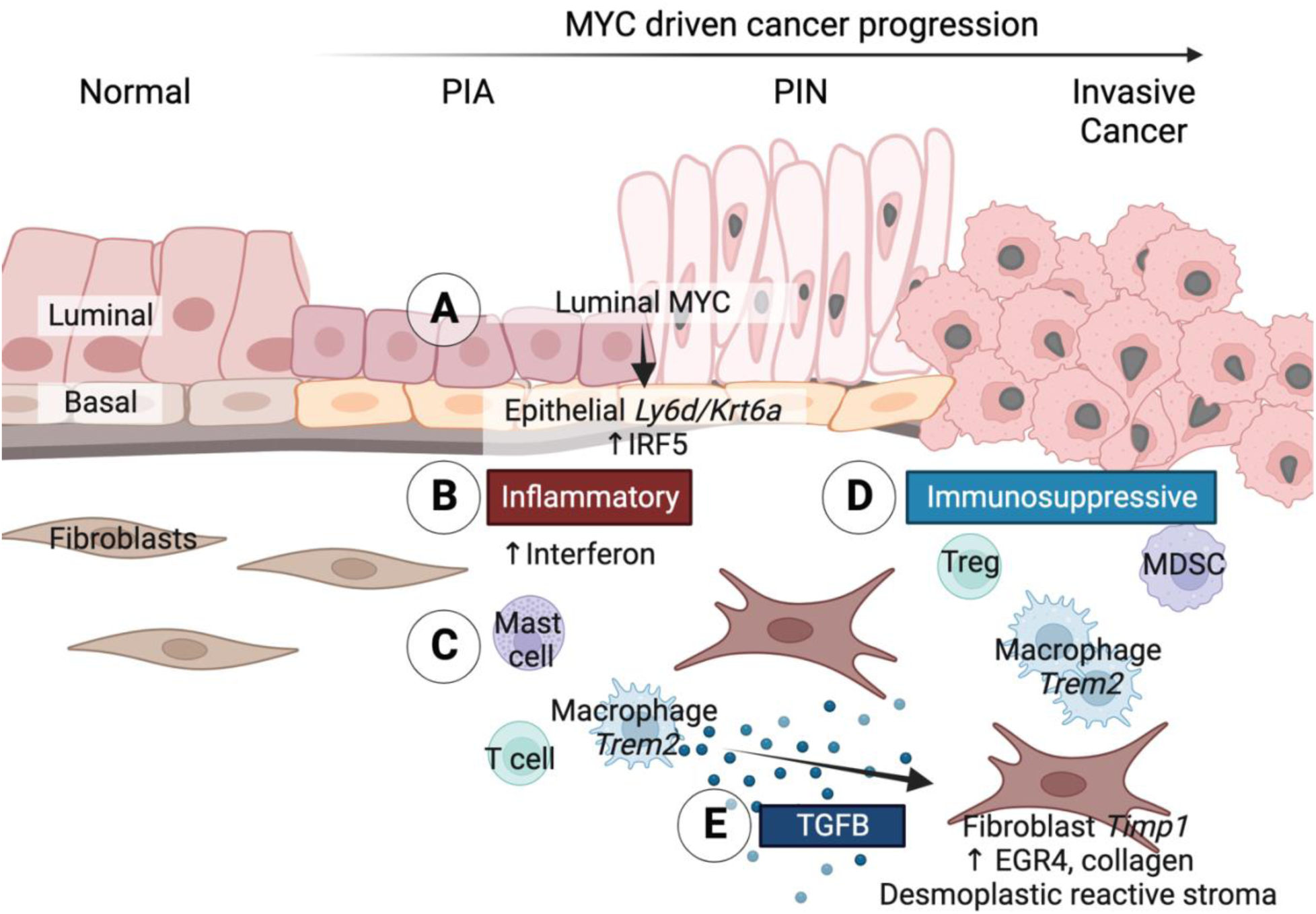
MYC activation in neoplastic cells reprograms the prostate TME. A) MYC activation in luminal epithelial cells leads to the induction of benign *Ly6d/Krt6a*-expressing epithelial population in the basal compartment of prostate glands. B) At the precursor stage, the TME is pro-inflammatory, with upregulated interferon signaling in both MYC-expressing luminal cells and Ly6d-expresing basal cells, and C) enrichment of various immune cells in the TME, including mast cells, T cells and *Trem2*-expressing macrophages. D) As MYC expressing luminal cells progress to invasive carcinoma, there is a pro-inflammatory to immunosuppressive switch with downregulation of inflammatory response pathways and IL6 JAK STAT3 signaling and enrichment of immunosuppressive cell types including regulatory T cells (Tregs), myeloid-derived suppressive cells (MDSCs), and *Trem2*-expressing macrophages. E) Secretion of TGFB by *Trem2*-expressing macrophages activates TGFB signaling and EGR4 transcription factor activity in fibroblasts, resulting in a desmoplastic CAF population expressing Timp1 and reactive-stroma associated ECM proteins such as collagen. Graphical abstract created in Biorender.com.

The TREM2+ macrophages express high levels of *Tgfb1*, which likely signals through TGFbeta receptors to induce Egr4 mediated expression of reactive-stroma-associated collagen genes and *Timp1* in a population of cancer associated fibroblasts that dominate the stromal microenvironment in the invasive cancer-enriched tissues. Consistent with these findings, a recent single-cell study of fibrosis in human liver tissues reported that TREM2 macrophages promoted fibrosis-associated changes in the stroma ^86^.

This cascade of tumor microenvironmental alterations to cell states in MYC driven murine prostate cancer had several key parallels to that in human prostate cancer. The initial pro-immunogenic programs induced in the Luminal MYC 1 cells in the mouse have strong parallels to the intermediate cell clusters in the human prostate, which also show activation of MYC signaling alongside strong activation of immunogenic programs. The Basal Ly6d/Krt6a cells seen in the precursor lesions have strong parallels to a LY6D/KRT6A cell population seen in the basal compartment of PIA, PIN, and intraductal cancer lesions in the human tissues. The Macrophage Trem2 and Fibroblast Timp1 populations also had strong parallels in the human cancer-enriched prostate tissues. These findings establish the human relevance of the tumor microenvironmental alterations accompanying MYC driven prostate cancer in the mouse model.

While these findings provide some fundamental insights into convergent aspects of prostate cancer cells and their microenvironment, multiple important questions are raised that will require significant additional studies. First, what are the specific mechanisms by which MYC activation leads to the initial induction of pro-immunogenic programs? Numerous prior studies have implicated multiple pathways connecting MYC activation to immunosupressive signaling, including suppression of interferon signaling gene expression and subsequent inflammatory response, downregulation of MHC class I antigen expression, and secretion of pro-tumorigenic cytokines (CCL2, CCL9, IL13, IL23, etc.) that recruit and polarize macrophages ^31, 87–90^. However, the precise mechanisms at play prior to immune suppression in the pro-inflammatory early stages of prostate carcinogenesis are not well established. Second, what specific mechanisms are responsible for the switch from the pro-immunogenic signals to an immunosuppressive state despite ongoing MYC activation and signaling? With the identification of the specific types of changes to the immune and stromal microenvironment associated with this switch in this study, it will be possible to uncover the mechanisms responsible for these changes in future studies.

These alterations induced by MYC driven prostate cancer may also represent convergent sequelae of other cancer drivers, perhaps via cross talk through MYC activation. For example, a recent scRNA-seq study in mouse models of prostate cancer driven by PTEN, RB1, and/or TP53 deletion reported upregulation of JAK/STAT inflammatory signaling was a key instigator of lineage plasticity in prostate cancer cells ^91^. Notably, adenocarcinoma cells in this model were also upregulated for inflammatory response, interferon signaling, and TNFA signaling via NF-κB.

In conclusion, MYC activation is a common denominator across all molecular subtypes of prostate cancer. MYC activation in luminal cells, directly and indirectly, reprograms the local TME, with widespread effects on neighboring epithelial, stromal, and immune cell types. These changes in the TME of prostate cancer may be informative for future therapeutic strategies. For example, there is an ongoing clinical trial with antibodies targeting TREM2 macrophages in cancer ^92^. Our study shows that one of the key changes in the TME of MYC-driven prostate cancer includes the enrichment of TREM2 macrophages with implications for driving an immunosuppressive and reactive stromal microenvironment.

## MATERIALS AND METHODS

### Human prostatectomy tissue punches

Prostate tissue specimens were collected from men diagnosed with primary prostate cancer undergoing radical prostatectomy at Johns Hopkins University (Table S1) and consented under IRBs (NA_00048544 and NA_00087094). Prostatectomies were sectioned fresh from apex to base, and fresh tissue samples were collected using an 8 mm punch biopsy tool from each zone of the prostate, as well as apparent sites of tumor (approximately 4 punches per subject). As described below, for each tissue punch, a portion of the punched tissue core was used to cut frozen sections for H&E staining, immunostaining (PIN4, ERG, PTEN, and MYC), and in situ hybridization for human MYC mRNA. The adjacent portion of the tissue punch was processed for scRNA-seq.

### Mouse prostate dissection

Wildtype C57BL/6J and FVB/NJ mice were purchased from Jackson Laboratory. The FVB Hi-Myc mice, FVB-Tg(ARR2/Pbsn-MYC)7Key, were purchased from NCI Mouse Repository. The laboratory of Brian Simons provided the C57BL/6J Hi-Myc mice (B6 Hi-Myc). The C57BL/6J Hi-Myc mice were generated by backcrossing the Hi-Hyc transgenic mice originally developed in FVB/NJ mice ^55^ with C57BL/6J for more than ten generations ^56^. Mice were maintained until they reached six months and ten months (+/- 6 weeks) of age. Individual prostate lobes were dissected, and scRNA-seq libraries for each lobe were prepared separately. Mice were euthanized using carbon dioxide asphyxiation, and the urogenital tract was removed and placed into a petri dish containing 50 ml of Hanks’ Balanced Salt Solution (HBSS, Gibco 14175-079). Under a dissection microscope, adipose tissues were removed from the urogenital tract to isolate the prostate. The four pairs of lobes (anterior, dorsal, lateral, and ventral lobes) were separated and dissected from the urethra by forceps. One pair was dissected in half for single-cell RNA-Seq, and the other half was formalin-fixed & paraffin-embedded (FFPE). The remaining lobe pair was frozen.

### Dissociation of mouse and human prostate tissues

Dissected tissues were processed as previously described ^59^. Briefly, tissues were minced with razor blades and digested in 0.25% Trypsin-EDTA (Gibco 25200-072) for 10 min at 37°C, followed by incubation for 2.5 hours at 37°C with gentle agitation in DMEM containing 10% FBS, 1 mg/mL Collagenase Type I (Gibco 17100-017), and 0.1 mg/mL of DNase I (Roche 10104159001). Digested tissues were centrifuged at 400 x g for 5 minutes, washed with HBSS, and further incubated in 0.25% Trypsin-EDTA for 10 min at 37°C. Cells were suspended in DMEM containing 10% FBS and 0.4 mg/mL of DNase I and filtered through a 40 μm cell strainer.

### Single-cell RNA-sequencing and data pre-processing

Libraries for scRNA-seq were prepared using the 10x Genomics Chromium Single Cell 3’ Library and Gel bead Kit V2 (CG00052_RevF) according to the manufacturer’s protocol for each dissected mouse prostate lobe and each human prostatectomy tissue punch. The cDNA libraries were sequenced (150 bp paired-end) on the Illumina HiSeqX platform. Each sequenced library was demultiplexed to FASTQ files using Cell Ranger (10x Genomics). For human tissues, Cell Ranger (version 3.2.0) count pipeline was used to align reads to the GRCh38 transcriptome and create a gene-by-cell count matrix. The human sequencing files can be accessed on NCBI dbGaP (In Progress). For mouse tissues, Cell Ranger (version 2.2.0) count pipeline was used to align reads to the mm10 mouse transcriptome and create a gene-by-cell count matrix. To align transgene expression, Kallisto quant (v0.46.0) was used to align the reads against human *MYC* and mouse *Myc* reference sequences. Samtools (v0.1.19) was used to extract alignments from BAM files for each sample. An in-house Perl script was used to map the alignments to the original FASTQ reads and create a feature-barcode count matrix. The Cell Ranger matrices were merged with the custom count matrices containing human *MYC* and mouse *Myc*. The mouse sequencing files can be accessed upon request. For both mouse and human datasets, Seurat (version 4.3.0) ^93^ was used to pre-process the data further. Cells with mitochondrial gene percentages >25% were removed.

### Dimensionality reduction & cell type identification

Normalization and variance stabilization of scRNA-seq data were performed using Seurat (version 4.3.0), specifically SCTransform (version 2) with gamma-Poisson generalized linear model fitting ^93–97^. For dimensionality reduction and clustering analysis, a principal component analysis was performed, and a range of 30 to 50 principal components was used to compute the Uniform Manifold Approximation & Projection (UMAP) dimensions and perform Louvain clustering at varying resolutions, ranging from 0.1 to 1.0. Differential gene expression analysis of previously characterized cell type-specific genes was used to identify the cell type for each cluster ^59, 98^. Mouse prostate data for aggregated 6-month WT and Hi-Myc scRNA-seq data were split by strain and normalized by SCTransform. To integrate datasets split by strain, 2000 anchor features were chosen, canonical correlation analysis was selected for dimensionality reduction with 20 dimensions used to specify the neighbor search space, and 200 neighbors (k-filter) were specified for anchor filtering. Mouse prostate scRNA-seq data for FVB/NJ dorsal and lateral lobes of 6-month and 10-month mice were split by individual samples, and each sample library was normalized by SCTransform. A WT sample and a Hi-Myc sample from 10-month-old mice were selected as references for integration.

### Differential gene expression analysis

To identify marker genes, differential gene expression analysis of all clusters was performed using Seurat (version 4.3.0) in merged datasets without integration. Genes considered in the analysis needed to be expressed in at least 25% of cells in a cluster with a minimum natural log fold change of 0.25. All outputs of this analysis are included as supplemental tables. The gene filtering parameters used to generate each heatmap or dot plot from differential gene expression analysis are described in the legends.

### Inferred copy number variation (inferCNV)

We applied computational methodologies to infer copy number variations (CNVs) in our prostatectomy scRNA-seq data. In this approach, transcripts from neighboring genes on chromosomes with consistent up- or down-regulated expression are inferred to have a genomic gain or loss, respectively ^99^. For each subject, luminal cells from the transition zone were used as a reference to infer CNVs in the luminal cells of peripheral zone tissue punches. To determine which genes were used for inferCNV analysis, genes with a mean count of 0.1 were selected. The residual expression intensities were denoised using default dynamic thresholding. A three-state (deletion, neutral, amplification) hidden Markov model-based method was used to predict CNVs. Cells were grouped based on similar CNV patterns using the “subclusters” analysis mode with the default Leiden method to partition hierarchical clustering trees. Subclusters with inferred CNVs previously reported in prostate cancer were identified and incorporated in the Seurat object as metadata to indicate predicated cancer cells on the UMAP.

### Pathway and Gene Set Enrichment Analysis (GSEA)

Gene sets from the Molecular Signatures Database (msigdb), including the Hallmark Collection, KEGG, Reactome, and Dang MYC Targets Up, were imported using the R package msigdbr (version 7.5.1) ^54, 100–104^. The Wilcoxon rank test was performed on genes differentially expressed between clusters using Presto (version 1.0) ^105^. The specific comparisons are indicated for each plot and figure legend. Genes were ranked by the statistical output, and GSEA was performed using the R package fgsea to determine normalized enrichment scores (NES) and associated adjusted p-values (version 1.22.0) ^106^. In the mouse scRNA-seq dataset, the Pearson correlation of human MYC transgene was determined for all genes. We used the Investigate Gene Sets tool, available on msigdb, to compute significant overlaps between genes that positively correlated with human *MYC* transgene expression and biological pathways annotated by KEGG and Reactome.

### Gene Signature Analysis

Gene signature scores were computed using UCell (version 2.0.1) to evaluate cluster-specific marker gene expression ^107^. Gene lists for gene signature analysis included the leading edge genes from GSEA of MYC V1 targets comparing luminal and cancer clusters, regulatory T cell markers (*IL2RA*, *FOXP3*, *CD4*, *IKZF2*, *CCR4*, *CTLA4*), and Luminal Timp1 cluster marker genes (Table S9). The resulting scores ranging from 0 to 1 for each cell are indicated in UMAP plots.

### Immunohistochemistry (IHC)

Frozen and formalin-fixed and paraffin-embedded (FFPE) tissues were immunostained using the Ventana Discovery Ultra Autostainer IHC (Roche Diagnostics, Basel, Switzerland). Tissue slides were steamed for 32 minutes in Cell Conditioning 1 (CC1) solution (Roche Diagnostics, Cat. No. 950-500) for antigen retrieval. For multiplex PIN4 cocktail staining, tissues were incubated with antibodies diluted 1:50, including CK903 (Enzo, Cat. No. ENZ-C34903) for 40 minutes, P63 (Biocare, Cat. No. CM163A) for 40 minutes, and AMACR (Zeta, Cat No. Z2001L) for 32 minutes. Chromogenic staining of target antigens was performed using the DISCOVERY anti-HQ HRP kit (903/p63 cocktail) and the Discovery anti-HQ-NP kit (AMACR). minutes at 1:50. Similarly, PTEN, ERG, CD3, F4/80, FOXp3 (Cell Signaling, Cat. No 12653) staining was performed on the Ventana Discovery Ultra Autostainer IHC. FFPE slides were steamed for 48 minutes in Cell Conditioning 1 (CC1) solution (Roche Diagnostics, Cat. No. 950-500) for antigen retrieval. Tissues were incubated with primary antibody using the following conditions, PTEN (Cell Signaling, Cat. No. 9188) diluted 1:100 with 1-hour incubation, ERG (Roche, Cat. No 6478450001) undiluted with 12 minutes of incubation, CD3 (FisherSci, Cat. No RM9107S) diluted 1:200 with 40 minutes of incubation, F4/80 (Cell Signaling, Ca. No 70076) diluted 1:200 with 36 minutes of incubation, and FOXp3 Cell Signaling (Cat. No 12653) diluted 1:250 with 1-hour incubation. Chromogenic staining of target antigens was performed using the DISCOVERY anti-HQ HRP kit.

### Chromogenic in situ hybridization (CISH)

To prepare frozen and FFPE tissue for CISH staining, slides were incubated for 30 minutes at 60°C and deparaffinized by incubating slides at room temperature (RT) for 10 minutes in xylene twice and then subsequently incubated in 100% ethanol twice and finally left to air dry. Hydrogen peroxide solution was added to the slides for 10 minutes at RT. Slides were steamed in 1 x RNAscope Target retrieval reagent at 100°C for 18 minutes, followed by protease plus digestion for 30 minutes at 40°C to allow target accessibility. The following probes were used for CISH staining, including Hs-MYC (RNAscope, Cat No. 311761) and Mm-Ly6d (RNAscope, Cat No. 532071) in mouse tissues. Probes were added to slides and incubated in the HybEZ TM Oven for 2 hours at 40°C. Frozen human prostatectomy tissue was cut into 5 μm sections and fixed in 10% neutral buffered formalin for 15 minutes at 4°C and dehydrated in alcohol. Hydrogen peroxide solution was added to the slides for 10 minutes at RT. Slides were steamed in 1 x RNAscope Target retrieval reagent at 100°C for 5 minutes, followed by protease IV digestion for 10 minutes at RT to allow target accessibility. Hs-KRT6A (RNAscope, Cat No. 520721) probe was added to slides and incubated in the HybEZ TM Oven for 2 hours at 40°C. Signal amplification and detection were performed using the RNAscope 2.5 HD Reagent kit-Brown (Cat.No 322300, ACD) according to the manufacturer’s protocol. The probe signal was detected with DAB. Slides were counter-stained with 50% Gill’s Hematoxylin for 2 min, rinsed in 0.02% ammonia water for 15 seconds, treated in 100% ethanol and xylene, and coverslipped with Cytoseal mounting medium.

### Cell communication network analysis

A computational strategy was used to construct inferred ligand-receptor-transcription factor cell communication networks among cell clusters in our mouse scRNA-seq datasets. Transcription factor activation scores were determined using the Single-Cell rEgulatory Network Inference and Clustering (SCENIC) pipeline ^108^. A Docker image provided by developers was used to run the Python implementation of SCENIC. The R-based package Domino was used to compile ligand-receptor interactions using the CellphoneDB repository ^109^, calculate the transcription factor activity enrichment for each cluster based on SCENIC-defined activation scores and correlate receptor expression and transcription factor scores ^110^. The parameters for constructing cell communication networks included a maximum of 10 transcription factors for each cluster with a minimum p-value of 1×10^-4^ and at correlation ≥ 0.25 for receptor expression and transcription factor activation.

## ACKNOWLEDGEMENTS

We thank the members of the Sidney Kimmel Comprehensive Cancer Center’s Experimental and Computational Genomics Core, supported by Cancer Center Support Grant P30CA006973, for support with the single-cell sequencing studies and data pre-processing and analysis. This work was supported in part by NIH/NCI grants P50CA058236, U01CA196390, P01CA247886, U54CA274370, DOD PCRP grant W81XWH-19-1-0450, and by the Prostate Cancer Foundation, The Allegheny Health Network Johns Hopkins Pilot Project Grant, The Patrick C. Walsh Fund, The Irving Hansen Foundation, The Commonwealth Foundation, and the Maryland Cigarette Restitution Fund.

