## Supplemental Information for "Convergent alterations in the tumor microenvironment of MYC-driven human and murine prostate cancer"

### SUPPLEMENTARY INFORMATION

#### Identifying benign and atrophic/intermediate epithelial cells

*GSTP1* and *LTF genes* known to be suppressed by promoter CpG island hypermethylation in prostate cancer cells and preneoplastic lesions [33–35], had reduced or absent expression in the prostate cancer clusters (Figure S1). In contrast, both *GSTP1* and *LTF* are known to be highly expressed in the luminal epithelial cells of proliferative inflammatory atrophy (PIA) lesions [18,35,41]. We observed that *LTF* and *GSTP1* showed high expression in a cluster of luminal cells, which we refer to as atrophic/intermediate. Consistent with upregulated inflammation, GSEA using the Hallmark Geneset Collection revealed that this atrophic/intermediate population was enriched for several immune pathways, including TNFA Signaling via NFKB, Inflammatory Response, and Interferon Gamma Response (Figure S7B), similar to a previously described luminal epithelial population with upregulated KLK3, KLK2, and KLK4 [40]. In addition to expressing genes that are upregulated in PIA, this cluster also expressed genes previously described in club cells and prostate-cancer-associated club/intermediate cells, including *PIGR*, *MMP7*, and *LCN2* [38,111]. Trajectory analysis of human prostate tissues suggested that these club/intermediate cells are luminal precursors [85].

#### Identifying neoplastic epithelial cells

Prostate cancer antigen 3 (*PCA3*), known to be upregulated in prostate cancer cells [36], is expressed in cancer clusters 1, 2, 5-7. Similarly, alpha-methyl acyl-CoA racemase (*AMACR*), known to be upregulated in prostate cancer and PIN[37,112], were expressed in cancer clusters 1-3, and 6-8 (Figure S1). The very low rate of AMACR-positive cells in cancer clusters 4 and 5 may be due to dropout [113]. PIN4 immunohistochemical stain shows heterogenous AMACR protein expression in subject 9, which largely comprises the cancer-4 cluster (Supplemental Figures S2A & D). The triple antibody cocktail PIN4 stains nuclear p63 and cytoplasmic high molecular weight cytokeratins brown in basal cells and cytoplasmic AMACR red [114].

#### Disease Progression in Hi-Myc mouse model of prostate cancer

The transgenic Hi-Myc mouse, originally generated in the FVB/NJ mouse strain, contains a construct of the human *MYC* oncogene with three androgen response elements upstream of the human transgene. Consequently, transgene expression is driven by androgen receptor activity. In this model, precursor PIN lesions can be observed as early as 2-4 weeks, with evidence of invasive adenocarcinoma beginning to develop between 3-6 months, transitioning to more extensive invasive adenocarcinoma in dorsal and lateral lobes after 6 months [55]. At approximately 8 months, tumors are visible upon dissection (Figure S8).

C57BL/6J Hi-Myc mice were previously generated by backcrossing Hi-Myc transgenic mice with C57BL/6J mice for more than ten generations [56]. It is known that strain differences can be observed in the Hi-Myc mouse model, with a significant delay in cancer progression in C57BL/6 compared to FVB/NJ [56,57]. We reasoned that by evaluating both strains, we could identify the convergent alterations in the neoplastic cells and their microenvironment in MYC-driven prostate carcinogenesis across this strain heterogeneity.

**Heterogeneous human MYC transgene expression across prostate lobes of Hi-Myc mice**

Chromogenic *in situ* hybridization (CISH) of human *MYC* confirmed robust transgene expression in invasive carcinoma and the absence of transgene expression in normal glands of WT mouse prostates (Figure 3B). Consistent with previous reports [51,55,57,58], we observed heterogeneous expression of human *MYC* transgene across the lobes of Hi-Myc mice in both strains, with the highest expression in the luminal cells of the ventral lobe, followed by lateral and dorsal lobes and the anterior lobe having the least expression (Figure 3C).

**Ly6d-expressing epithelial cells enriched in Hi-Myc mouse prostate**

While *Ly6d* has been previously described as a urethral marker in WT mice [38], the urethra was not included in the dissection and scRNA-seq library preparation of the mouse prostates in this study, suggesting that these Ly6d-positive, non-urethral cell clusters selectively emerged in the context of the Hi-Myc mice. Notably, these Ly6d-positive cells express the prostate-specific marker *Ceacam1* and do not robustly express the peri-urethral specific marker *Aqp3*, previously described by Crowley et al. [60] (Supplemental Figure S10A). The luminal Ly6d cluster, but not the Basal Ly6d cluster, also expressed prostate stem cell antigen (*Psca*), suggesting that this cluster includes the previously described rare luminal population with high progenitor potential [60,115–118]. A previous scRNA-seq study of Hi-Myc mice reported that the *MYC* transgene did not transcriptionally reprogram this luminal progenitor population [51]; however, these studies were performed in 12-week Hi-Myc mice when the prostate is comprised primarily of precursor lesions, in contrast to our dataset of 6-month-old Hi-Myc mice, where both PIN and invasive carcinoma are present (Figure S8).

A recent scRNA-seq study of a Pten knockout mouse model of prostate cancer also reported a unique epithelial cluster, which they described as an intermediate epithelial population expressing keratin 4 (*Krt4)*, tumor-associated calcium signal transducer 2 (*Tacstd2*), and protein phosphatase 1 regulatory inhibitor subunit 1B (*Ppp1r1b*) but lacking *Psca* expression, making it distinct from the previously reported *Psca* expressing progenitor population [119]. While sharing some similar gene expression with the luminal Ly6d cluster, the expression of Psca suggests that the luminal Ly6d cluster is distinct from the previously described intermediate cells observed in the *Pten* loss mouse model of prostate cancer (Figure S10A). Gene expression analysis revealed that Ly6d-expressing luminal cells had several genes that were differentially expressed between the two clusters (Figure S10A); for example, *Psca*, peptidoglycan recognition protein 1 (*Pglyrp1*), galectin 7 (*Lgals7*), and carboxypeptidase E (*Cpe*).

**A subset of MYC-expressing epithelial cells express beta-defensins**

Beta defensins are antibiotic peptides known to be expressed in epithelial cells and are chemotactic for various immune cell types [120]. In particular, beta defensins have been shown to be a potent chemoattractant for macrophages [121–123]. We observed that the Hi-Myc mouse prostate was significantly enriched for a luminal cluster, referred to as Luminal MYC 2, expressing beta defensins *Defb29* and, to a lesser extent *Defb28* (Figure C, Supplemental Figure S10C, Table S5). In humans, we observed that a subset of cells within the basal and intermediate/atrophic clusters in our prostatectomy samples expressed the beta defensin *DEFB1* (Supplemental Figure S10D).

**Inferring paracrine interactions between MYC-expressing luminal cells and neighboring Ly6d-expressing basal cells**

To explore potential paracrine interactions between MYC-expressing luminal cells (Luminal Timp1 cluster) and *Ly6d*-expressing basal cells, we employed the computational tool, Domino, to construct ligand-receptor-transcription factor cell communication networks from scRNA-seq data [110]. Many of the top expressing ligands targeting the Basal Ly6d cluster are expressed by the Luminal Timp1 cluster, including interferon kappa (Ifnk), fibrinogen beta chain (Fgb), Wnt family member 5a (Wnt5a), and bone morphogenetic protein 3 (Bmp3) (Figure 5B). Transcription factor activity scores derived from SCENIC, a  computational method to examine gene regulatory networks in scRNA-seq datasets [108], identified upregulated activity of early growth response 2 (EGR2), SRY-Box Transcription Factor 4 (SOX4), transformation-related protein 63 (TRP63), forkhead box A2 (FOXA2), interferon regulatory factor 5 (IRF5), MAF BZIP transcription factor B (MAFB), retinoic acid receptor gamma (RARG), MYB proto-oncogene like

1 (MYBL1), oligodendrocyte transcription factor 1 (OLIG1), and zinc finger protein 958 (ZFP958) in the Basal Ly6d cluster.

Coupling ligand-receptor pair interactions with transcription factor activity, we observed a positive correlation between interferon regulatory factor 5 (IRF5) activity with the expression of multiple class II cytokine receptors, including interferon lambda receptor 1 (*Ifnlr1*), interleukin 10 receptor subunit beta (*Il10rb*), and the type 1 interferon receptor subunits, interferon alpha and beta receptor subunit 1 (*Ifnar1*) and interferon alpha and beta receptor subunit 2 (*Ifnar2*) (Figures 5B-C). Similar to other type 1 interferons, such as IFN alpha and IFN beta, IFNK binds to the type 1 interferon receptor [64,65]. In general, *IRF5* is expressed in lymphoid cells; however, *IRF5* expressed in other cell types, including epithelial cells [124,125]. Furthermore, the downstream targets of IRF5, as defined by SCENIC regulatory networks, were upregulated in the Basal Ly6d cluster (Figures 5D), including gelsolin (*Gsn*), TIMP metallopeptidase inhibitor 2 (*Tmp2*), out at first homolog (*Oaf*), CD47 molecule (*Cd47*), and syntaxin 1A (*Stx1*).

**Strain-specific differences in the myeloid population of Hi-Myc mice**

Further examination of the immune population in Hi-Myc mice (6 months) revealed three macrophage subclusters (Figures S12A-C) including a *Trem2* expressing cluster (Figure S12D-E), a *Cd72* expressing cluster (Macrophages Cd72) enriched in C57BL6/J mice, and a *Cxcl2* expressing cluster (Macrophages Cxcl2) enriched in FVB/NJ mice (Figures S12B, S12F). Notably, others have suggested differences in a myeloid population may be responsible for the delayed tumor progression observed in C57BL/6 mice compared to FVB/NJ Hi-Myc mice [57]. Regardless of strain-specific differences in the macrophage population, the prostate TME of Hi-Myc mice was significantly enriched for macrophages, T cells, and mast cells (Figure S12G). In particular, the *Trem2*-expressing macrophages were strain agnostic (Figures S12E-G).

**Fibroblast Timp1 cluster is enriched for CAF gene signatures**

We performed a gene signature analysis of our stromal clusters using the previously reported pan-CAF gene signatures [73]. Compared to other stromal clusters, the Fibroblast Timp1 cluster was enriched for the demoplastic pan-dCAF, inflammatory pan-iCAF, and proliferating pan-pCAF gene signatures (Supplemental Figure S17A). The range of low and high gene signature scores in the Fibroblast Timp1 cluster suggested that there may be a mixture of pan-dCAF, pan-iCAF, and pan-pCAF subpopulations. Evaluating the pan-CAF scores for individual fibroblast cells revealed that many cells had high pan-CAF scores ( $> 0.3$) for more than one pan-CAF gene signature (Supplemental Figures S17B-C).

### SUPPLEMENTARY FIGURES

**Figure S1. Epithelial composition of benign- and cancer-enriched prostatectomy tissues.** A) Feature plot shows the distribution of cancer-associated gene expression. *GSTP1* and *LTF* are down-regulated in prostate cancer clusters 1-8. *PCA3* is up-regulated in cancer clusters 1-2 and 5-7. *AMACR* is up-regulated in cancer clusters 1-3 and 6-8. *ERG* is up-regulated in cancer clusters 2, 4, and 5. Cancer-4 cluster cells show a complete loss of *PTEN* expression. B) Heatmap shows *SPINK1* upregulated in *ERG*-negative cancer clusters 1, 3, 6-8. C) Heatmap of top-upregulated and down-regulated genes by fold change for each epithelial cluster with unsupervised hierarchical clustering. Refer to Table S3 for a complete list of differentially expressed genes. D) Dotplot of top differentially expressed genes for each epithelial cluster. For each cluster, selected genes were expressed in less than 20% of other cells and ranked by adjusted p-value, followed by log fold change.

**Figure S2. Fresh Frozen Prostatectomy tissue immunostained for PIN4, ERG, and PTEN.** A) UMAPs highlighting ERG-negative/PTEN-positive cancer clusters 1, 3, and 6-8 (left panel), ERG-positive/PTEN-positive cancer clusters 2 and 5 (middle panel), and ERG-positive/PTEN-negative cancer cluster 4 (right panel). B) ERG negative/PTEN positive cancer regions (highlighted by arrows) from subject 7, consistent with cancer cluster 6. C) ERG positive/PTEN positive cancer regions (highlighted by arrows) from Subject 5, consistent with cancer cluster 2. D) ERG positive/PTEN negative cancer regions (highlighted by arrows) from Subject 9, consistent with cancer cluster 4. PIN4 cocktail staining (top row) is comprised of IHC for tumor protein P63 (TP63; brown), high molecular weight cytokeratin expressed in basal epithelial cells (brown staining), and alpha-methylacyl-CoA racemase (AMACR; red staining), which is over-expressed in carcinoma and prostatic intraepithelial neoplasia (PIN).

**Figure S3. Inferred copy number variation (CNV) for subjects 1, 2, and 4.** Heatmap from inferCNV analysis indicating chromosome regions with inferred CNV gain (red) and loss (blue) from representative subjects: A) Subject 1, B) Subject 2, and C) Subject 4. InferCNV analysis did not identify any known prostate cancer-associated CNVs in Subject 3. The “subgroup” category is derived from the unsupervised hierarchical clustering of individual cells and indicates copy number clusters. Arrows indicate which subgroups were determined to be consistent with cancer cells. The group category indicates tissue sample, peripheral zone benign-enriched (PZ), PZ cancer-enriched (CA), or central zone (CZ). Not shown is the reference sample from the transition zone (TZ). The group cell cluster represents the luminal epithelial group from dimensionality reduction and clustering analysis performed for each subject.

**Figure S4. Inferred CNVs for subjects 5, 7, and 8.** Heatmap from inferCNV analysis indicating chromosome regions with inferred CNV gain (red) and loss (blue) from A) Subject 5, B) Subject 7, and C) Subject 8. InferCNV analysis did not identify any known prostate cancer-associated CNVs in Subject 6.

**Figure S5. Inferred CNVs for subjects 9 and 10.** Heatmap from inferCNV analysis of cancer cells indicating chromosome regions with inferred CNV gain (red) and loss (blue) from A) Subject 9 and B) Subject 10.

**Figure S6. MYC is activated across all molecular subtypes of primary prostate cancer.** Heatmap of Hallmark MYC targets V1 gene expression in TCGA primary prostate cancer dataset. Rug chart annotations include androgen receptor (AR) activity score, clinical Gleason score, and molecular subtype. Additionally, for each cancer sample, the gene set enrichment analysis (GSEA) normalized enrichment scores (NES) for Dang MYC targets up, Hallmark MYC targets V2, and Hallmark MYC targets V1 were determined and color-scaled and displayed as rug charts on the top of the heatmap.

**Figure S7. Gene expression programs and MYC transgene activity distinguish distinct epithelial cell clusters.** Plot showing NES of top 20 Hallmark gene sets by adjusted p-value, comparing A) basal and luminal clusters, B) atrophic/intermediate and luminal clusters, and C) aggregated cancer groups and atrophic/intermediate clusters by GSEA. UMAPs of D) epithelial groups considered for GSEA, E) MYC expression, and F) signature score derived from MYC target genes. The leading edge genes of MYC targets V1 comparing cancer and luminal clusters, were used to derive the signature score. G) Representative examples of hematoxylin and eosin (H&E) staining of fresh frozen peripheral zone tissue punches that were benign-enriched (top panels, normal gland magnified) or cancer-enriched (bottom panels, invasive carcinoma magnified). PIN4 staining in the acini of normal glands shows TP63 and high molecular weight cytokeratin expression (brown) in basal epithelial cells lining the perimeter of normal luminal epithelial cells lacking AMACR (red) in the acini of normal glands.

Immunohistochemical staining shows the absence of MYC in normal luminal cells, while a subset of basal cells is positive for MYC. In contrast, PIN4 and MYC immunostaining in adenocarcinoma shows AMACR-expressing prostate cancer cells are positive for MYC.

**Figure S8. Time course of developing PIN, cribriform PIN/CIS, and invasive cancer in the Hi-MYC prostate cancer model, with key transitions occurring at 6 months and 8-10 months.** A) Plot showing the progression of disease in Hi-Myc prostate. Precursor PIN lesions develop as early as 2-4 weeks, followed by invasive carcinomas between 3-6 months. At approximately 8 months, invasive carcinoma is extensive, and tumors are visible upon dissection. B) Representative images of WT mouse prostate and Hi-Myc prostates at 6 months and > 8 months. Anterior, dorsal, lateral, and ventral lobes are indicated. C) Representative H&E images showing the spectrum of glands and lesions (precursor and carcinoma) observed in the prostate of 6-month Hi-Myc mice.

**Figure S9. Dimensionality reduction of unintegrated scRNA-seq mouse libraries.** A) UMAPs of mouse prostate cells grouped by cell type, strain (C57BL/6J and FVB/NJ), lobe (anterior, dorsal, lateral, ventral), and genotype (WT and Hi-Myc). B) UMAPs showing the expression of endogenous mouse *Myc*, the human *MYC* transgene, and ribosomal protein S5 (*Rps5*) – a downstream target of MYC.

**Figure S10. *Ly6d* and *Krt6a* are expressed in a subset of basal and luminal cells.** A) Dotplot of top 15 differentially expressed genes by adjusted p-value and log fold change for luminal and basal Ly6d/Krt6a expressing clusters. B) UMAP of mouse epithelial clusters (left panel), expression of beta defensin *Defb29* (middle panel), and co-expression of *Ly6d* and *Krt6a* (right panel). C) UMAP of human epithelial clusters (left panel), expression of beta defensin *DEFB1* (middle panel), and co-expression of *LY6D* and *KRT6A* (right panel).

**Figure S11. *KRT6A* is expressed in a subset of epithelial cells in the human prostate gland.** Example of *KRT6A* CISH staining primarily basal compartment of A) PIN (blue arrows) from subject 2 and B) PIA from subject 9 (black arrows) in human prostate from frozen tissue sections.

**Figure S12. Interferon response factor 5 (*Irf5*) is expressed in basal and luminal clusters expressing *Ly6d* and *Krt6a*.** UMAP of mouse epithelial clusters (left panel) and expression of the transcription factor *Irf5* (right panel).

**Figure S13. *Trem2*-expressing macrophages are enriched in Hi-Myc mice in both C57BL6/J and FVB/NJ strains.** A) Immune cell clusters were subsetted from 6-month Hi-Myc mouse prostates from C57BL6/J and FVB/NJ strains, and dimensionality reduction (UMAP) and clustering analysis were repeated. B) UMAP of immune cells marked by strain. C) Heatmap of top marker genes expressed in each immune cluster, separated by strain. D) UMAP shows *Trem2* expression in macrophages. E) Violin plot showing *Trem2* expression for each immune cluster, separated by strain. Scatter plots show the proportion of immune cell populations for each sample by F) strain or g) genotype. Each point represents a sample colored by genotype or age. Bar represents the mean expression of samples for each group. Statistics were generated by comparing Hi-Myc samples by strain or genotype using RAISIN's cell proportions test.

**Figure S14. Cell intrinsic changes in the Hi-Myc luminal population during cancer progression.** A) UMAP of FVB/NJ prostates from 6-month and 10-month Hi-Myc and age-matched dorsal and lateral lobes. Samples were integrated by age. B) Heatmap of cell type marker genes show cluster-specific expression. C) Pathways showing the greatest difference in GSEA NES in MYC-expressing luminal cells at 6 months (Luminal Timp1 cluster vs. Luminal cluster) and 10 months (Luminal Timp1 cluster 10 months vs. 6 months).

**Figure S15. Immunosuppressive cells in Hi-Myc prostate.** Immunostaining for FOXP3 in tissues from 6 and 10-month Hi-Myc and age-matched WT prostates.

**Figure S16. Stromal clusters in mouse and human prostates.** A) UMAP of stromal cells from FVB/NJ prostates from 6-month and 10-month Hi-Myc and age-matched dorsal and lateral lobes. Samples were integrated by age. B) Heatmap of cell type marker genes show cluster-specific expression in mouse stromal cells. C) UMAP of human stromal clusters from prostatectomy samples and D) associated heatmap showing cluster-specific expression of stromal genes.

**Figure S17. Timp1-expressing fibroblasts enriched for cancer-associated fibroblast gene signatures (Pan-CAFs).** A) Violin plots showing gene signature scores of the pan-CAF gene signatures across across stromal clusters from FVB/NJ prostates from 6-month and 10-month Hi-Myc. B) Heatmap of fibroblast cells and pan-CAF gene signature scores. C) UMAPs showing of fibroblast clusters (top panel) and cells with high (gene signature scores > 0.3, bottom panel).

**Figure S18. *Timp1*-expressing fibroblasts are enriched in Hi-Myc mice in C57BL6/J and FVB/NJ strains.** A) Stroma cell clusters were subsetted from 6-month Hi-Myc mouse prostates from C57BL6/J and FVB/NJ strains, and dimensionality reduction (UMAP) and clustering analysis were repeated (left panel). UMAPs of stromal cells marked by strain (middle panel) and genotype (right panel). B) Heatmap of cell type marker genes show cluster-specific expression in mouse stromal cells. Scatter plots show the proportion of stroma cell populations for each sample by C) strain or D) genotype. Each point represents a sample colored by genotype or age. Bar represents the mean expression of samples for each group. Statistics were generated by comparing Hi-Myc samples by strain or genotype using RAISIN's cell proportions test.

### SUPPLEMENTARY TABLES

**Table S1. Prostatectomy sample summary of cancer-enriched and benign-enriched tissue punches processed for scRNA-seq and histology.** Cell counts indicate the number of cells in the scRNA-seq dataset.

**Table S2. List of genes differentially expressed for each cell type cluster in prostatectomy samples.** Pct.1 indicates % of cells that the gene is detected in the cluster. At least 10% of cells must express the gene. Pct.2 indicates % of all other cells expressing the gene. The adjusted p-value is based on Bonferroni correction. The list of genes considered has a Bonferroni corrected p-value < 0.05 and log-fold change expression > 0.25 for a cluster of interest.

**Table S3. List of genes differentially expressed for each epithelial cluster in prostatectomy samples.**

**Table S4. Results from gene set enrichment analysis (GSEA) comparing cancer and luminal clusters from prostatectomy samples using Hallmark gene set collection.**

**Table S5. List of genes differentially expressed for each epithelial cluster in 6-month Hi-Myc and age-matched mouse prostates.** Samples included in this analysis include anterior, dorsal, lateral, and ventral lobes from FVB/NJ and C57BL6/J mouse strains.

**Table S6. List of genes differentially expressed for each cell type cluster in 6-month and 10-month Hi-Myc and age-matched mouse prostates.** Samples included in this analysis include dorsal and lateral lobes from FVB/NJ mouse strain.

**Table S7. List of genes differentially expressed for each stromal cluster in 6-month and 10-month Hi-Myc and age-matched mouse prostates.** Samples included in this analysis include dorsal and lateral lobes from FVB/NJ mouse strain.

**Table S8. List of genes differentially expressed for each stromal cluster in 6-month Hi-Myc and age-matched mouse prostates.** Samples included in this analysis include anterior, dorsal, lateral, and ventral lobes from FVB/NJ and C57BL6/J mouse strains.

**Table S9. The fibroblast Timp1 cluster gene signature.** The mouse (MGI) and human (HGNC) gene symbols are listed.

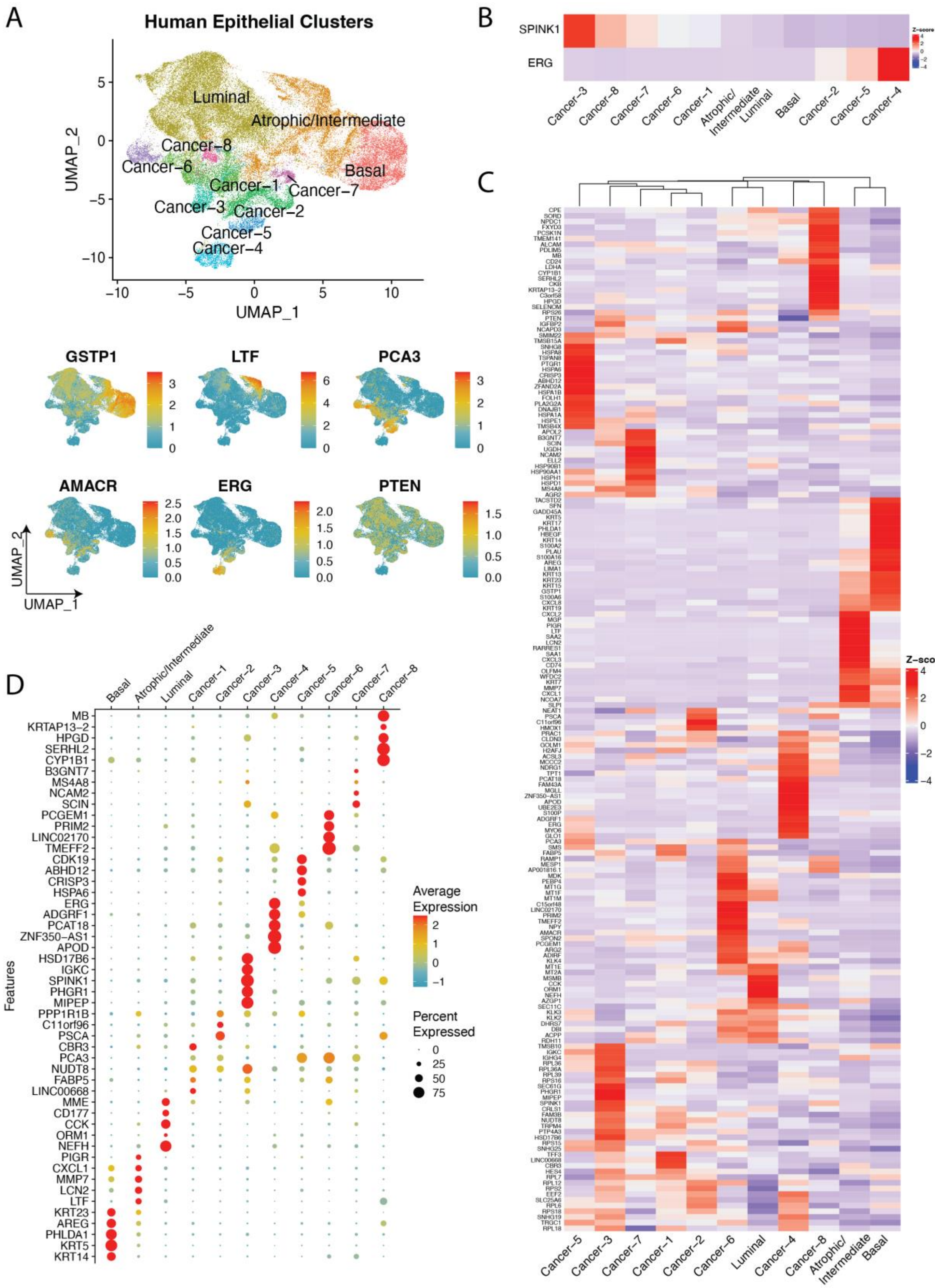
A
Human Epithelial Clusters
Luminal
Atrophic/Intermediate
Cancer−8
Cancer−6
Cancer−1
Basal
Cancer−7
Cancer−3
Cancer−2
Cancer−5
Cancer−4
UMAP_1
UMAP_2
GSTP1
LTF
PCA3
AMACR
ERG
PTEN
B
SPINK1
ERG
Z-score
Cancer−3
Cancer−8
Cancer−7
Cancer−6
Cancer−1
Atrophic/Intermediate
Luminal
Basal
Cancer−2
Cancer−5
Cancer−4
C
Z-score
Cancer−5
Cancer−3
Cancer−7
Cancer−1
Cancer−2
Cancer−6
Luminal
Cancer−4
Cancer−8
Atrophic/Intermediate
Basal
D
Basal
Atrophic/Intermediate
Luminal
Cancer−1
Cancer−2
Cancer−3
Cancer−4
Cancer−5
Cancer−6
Cancer−7
Cancer−8
Features
MB
KRTAP13−2
HPGD
SERHL2
CYP1B1
B3GNT7
MS4A8
NCAM2
SCIN
PCGEM1
PRIM2
LINC02170
TMEFF2
CDK19
ABHD12
CRISP3
HSPA6
ERG
ADGRF1
PCAT18
ZNF350−AS1
APOD
HSD17B6
IGKC
SPINK1
PHGR1
MIPEP
PPP1R1B
C11orf96
PSCA
CBR3
PCA3
NUDT8
FABP5
LINC00668
MME
CD177
CCK
ORM1
NEFH
PIGR
CXCL1
MMP7
LCN2
LTF
KRT23
AREG
PHLDA1
KRT5
KRT14
Average Expression
Percent Expressed
0
25
50
75

**Figure S1**

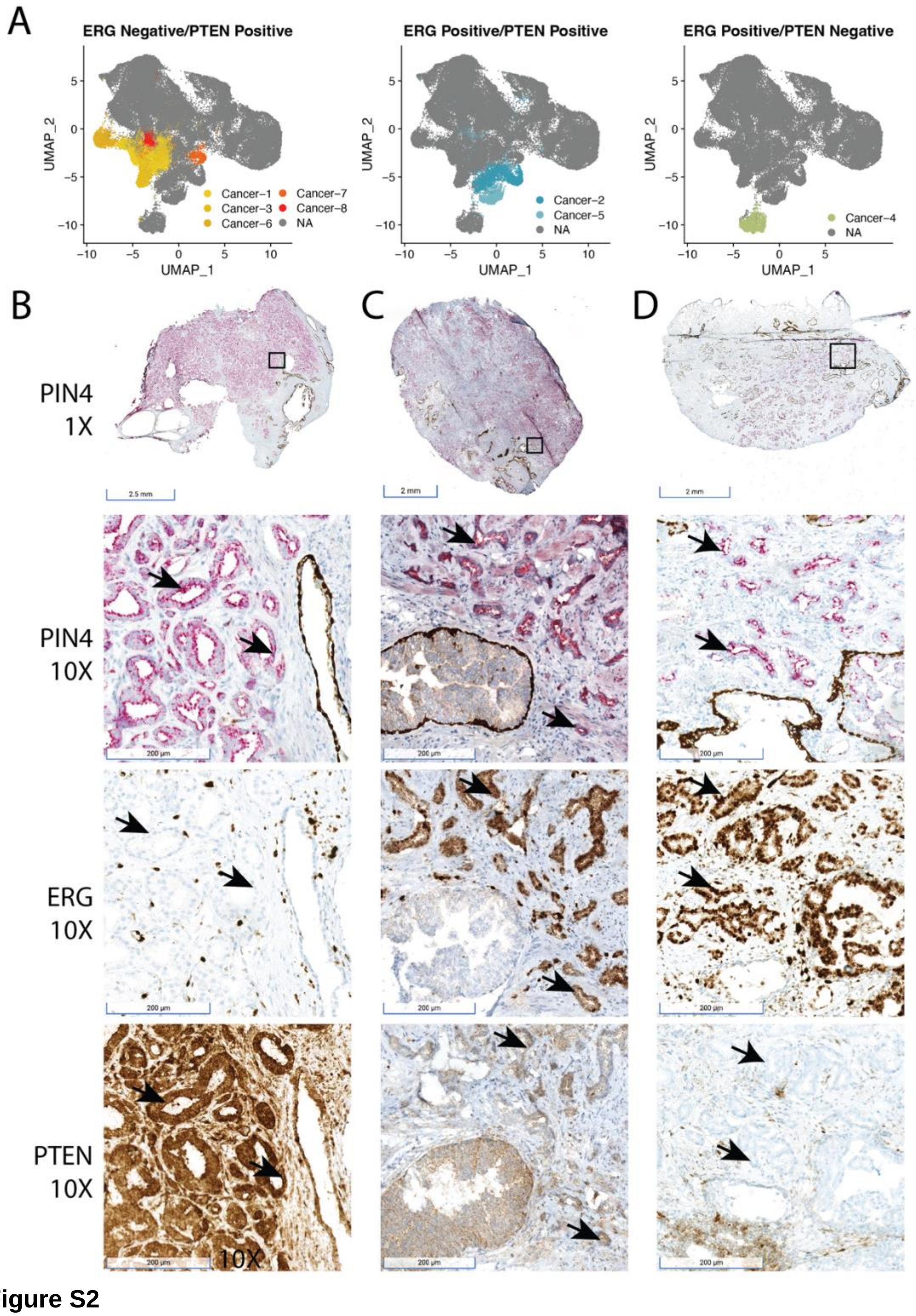
A
ERG Negative/PTEN Positive
ERG Positive/PTEN Positive
ERG Positive/PTEN Negative
UMAP_2
UMAP_1
Cancer−1
Cancer−3
Cancer−6
Cancer−7
Cancer−8
NA
Cancer−2
Cancer−5
Cancer−4
B
C
D
PIN4
1X
2.5 mm
2 mm
PIN4
10X
ERG
10X
PTEN
10X
200 µm
10X

**Figure S2**

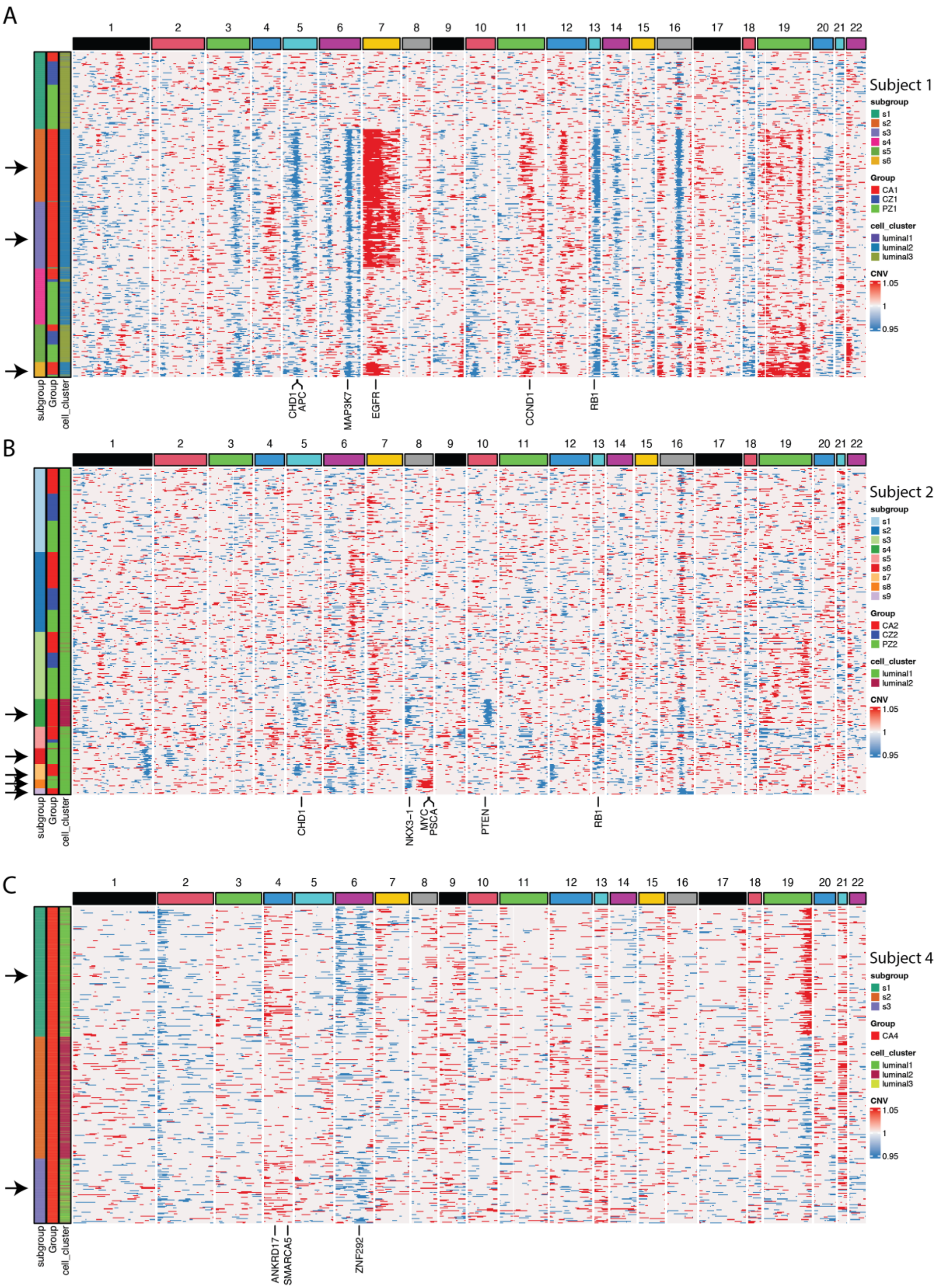

A
1 2 3 4 5 6 7 8 9 10 11 12 13 14 15 16 17 18 19 20 21 22
Subject 1
subgroup
s1
s2
s3
s4
s5
s6
Group
CA1
CZ1
PZ1
cell_cluster
luminal1
luminal2
luminal3
CNV
1.05
1
0.95
subgroup
Group
cell_cluster
CHD1
APC
MAP3K7
EGFR
CCND1
RB1
B
1 2 3 4 5 6 7 8 9 10 11 12 13 14 15 16 17 18 19 20 21 22
Subject 2
subgroup
s1
s2
s3
s4
s5
s6
s7
s8
s9
Group
CA2
CZ2
PZ2
cell_cluster
luminal1
luminal2
CNV
1.05
1
0.95
subgroup
Group
cell_cluster
CHD1
NKX3-1
MYC
PSCA
PTEN
RB1
C
1 2 3 4 5 6 7 8 9 10 11 12 13 14 15 16 17 18 19 20 21 22
Subject 4
subgroup
s1
s2
s3
Group
CA4
cell_cluster
luminal1
luminal2
luminal3
CNV
1.05
1
0.95
subgroup
Group
cell_cluster
ANKRD17
SMARCA5
ZNF292


**Figure S3**

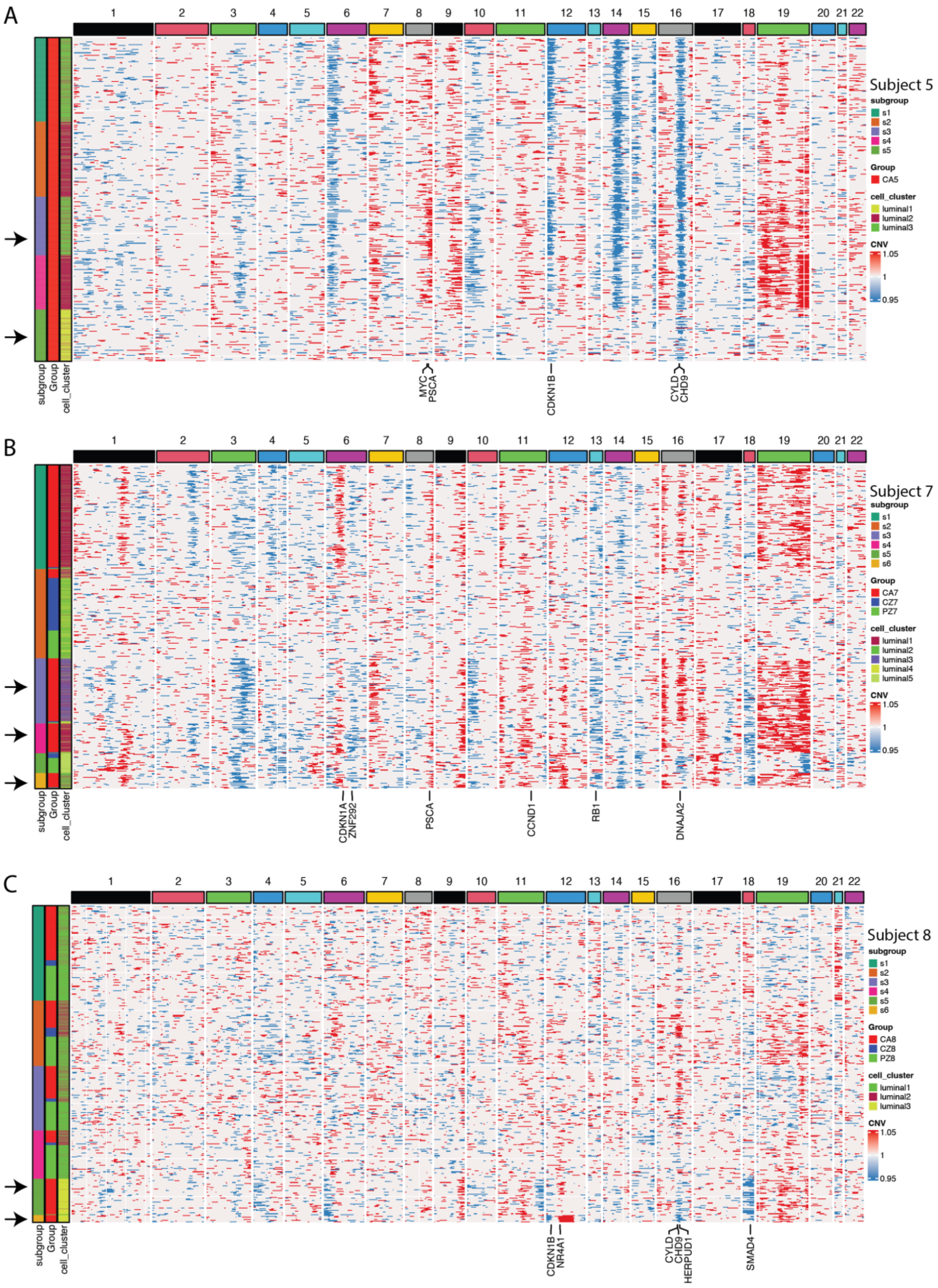

A
1 2 3 4 5 6 7 8 9 10 11 12 13 14 15 16 17 18 19 20 21 22
Subject 5
subgroup
s1
s2
s3
s4
s5
Group
CA5
cell_cluster
luminal1
luminal2
luminal3
CNV
1.05
1
0.95
subgroup
Group
cell_cluster
MYC
PSCA
CDKN1B
CYLD
CHD9
B
1 2 3 4 5 6 7 8 9 10 11 12 13 14 15 16 17 18 19 20 21 22
Subject 7
subgroup
s1
s2
s3
s4
s5
s6
Group
CA7
CZ7
PZ7
cell_cluster
luminal1
luminal2
luminal3
luminal4
luminal5
CNV
1.05
1
0.95
subgroup
Group
cell_cluster
CDKN1A
ZNF292
PSCA
CCND1
RB1
DNAJA2
C
1 2 3 4 5 6 7 8 9 10 11 12 13 14 15 16 17 18 19 20 21 22
Subject 8
subgroup
s1
s2
s3
s4
s5
s6
Group
CA8
CZ8
PZ8
cell_cluster
luminal1
luminal2
luminal3
CNV
1.05
1
0.95
subgroup
Group
cell_cluster
CDKN1B
NR4A1
CYLD
CHD9
HERPUD1
SMAD4


**Figure S4**

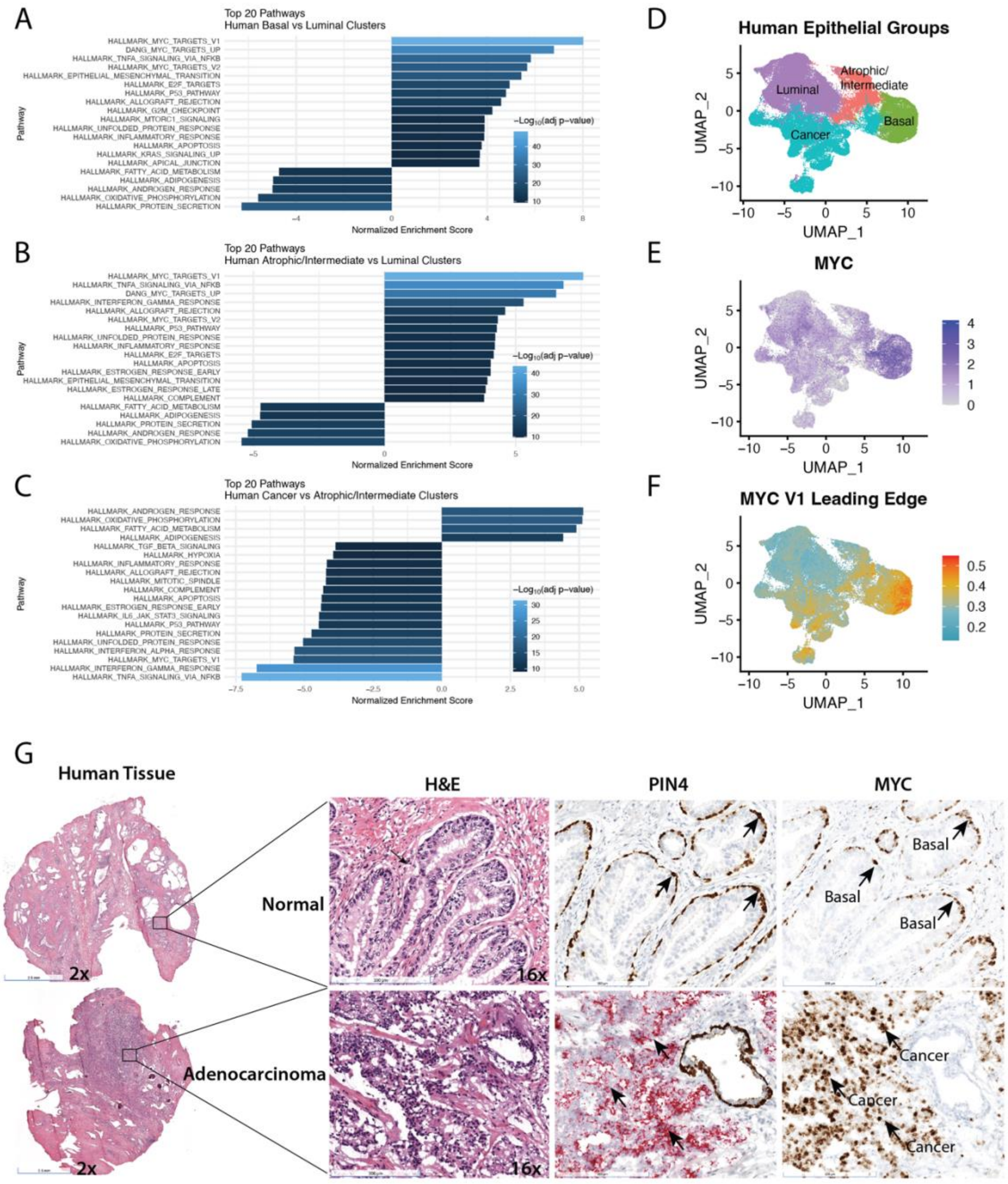

A
Top 20 Pathways
Human Basal vs Luminal Clusters
B
Top 20 Pathways
Human Atrophic/Intermediate vs Luminal Clusters
C
Top 20 Pathways
Human Cancer vs Atrophic/Intermediate Clusters
Normalized Enrichment Score
Pathway
D
Human Epithelial Groups
Luminal
Atrophic/
Intermediate
Basal
Cancer
UMAP_1
UMAP_2
E
MYC
F
MYC V1 Leading Edge
G
Human Tissue
H&E
PIN4
MYC
Normal
Adenocarcinoma
Basal
Cancer
2x
16x


Figure S5

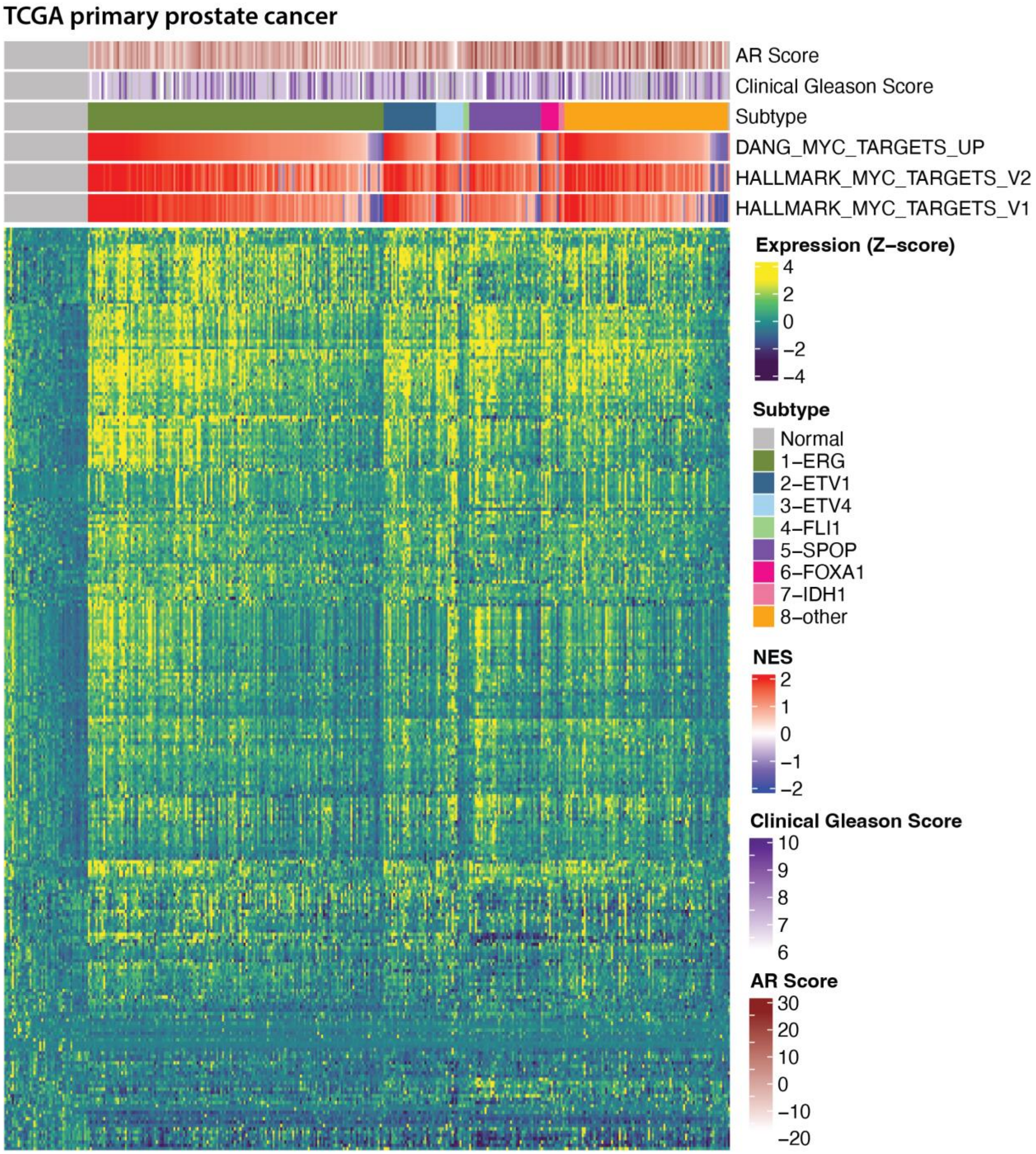
TCGA primary prostate cancer
AR Score
Clinical Gleason Score
Subtype
DANG_MYC_TARGETS_UP
HALLMARK_MYC_TARGETS_V2
HALLMARK_MYC_TARGETS_V1
Expression (Z−score)
4
2
0
−2
−4
Subtype
Normal
1−ERG
2−ETV1
3−ETV4
4−FLI1
5−SPOP
6−FOXA1
7−IDH1
8−other
NES
2
1
0
−1
−2
Clinical Gleason Score
10
9
8
7
6
AR Score
30
20
10
0
−10
−20

**Figure S6**

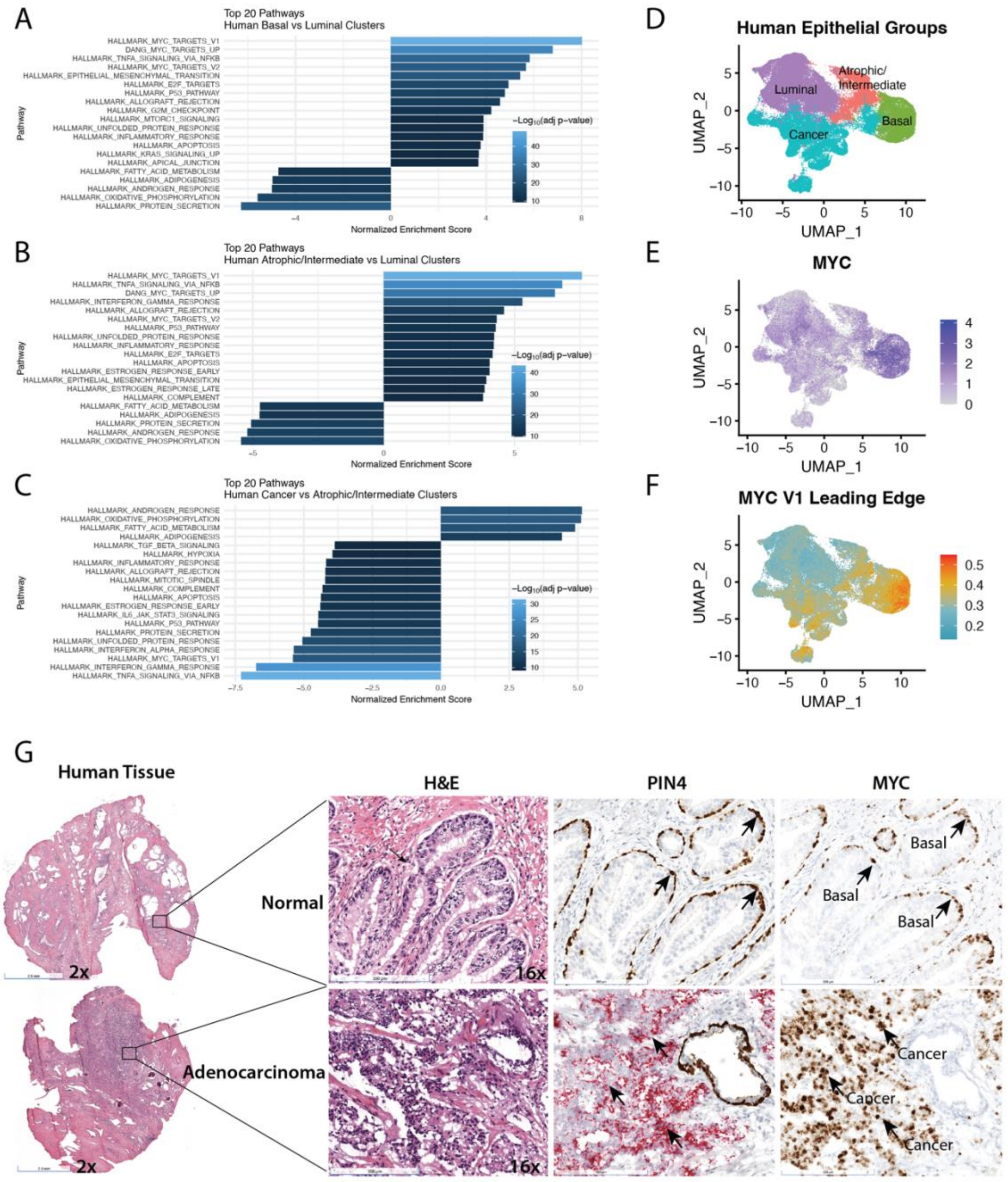
A
Top 20 Pathways
Human Basal vs Luminal Clusters
HALLMARK_MYC_TARGETS_V1
DANG_MYC_TARGETS_UP
HALLMARK_TNFA_SIGNALING_VIA_NFKB
HALLMARK_MYC_TARGETS_V2
HALLMARK_EPITHELIAL_MESENCHYMAL_TRANSITION
HALLMARK_E2F_TARGETS
HALLMARK_P53_PATHWAY
HALLMARK_ALLOGRAFT_REJECTION
HALLMARK_G2M_CHECKPOINT
HALLMARK_MTORC1_SIGNALING
HALLMARK_UNFOLDED_PROTEIN_RESPONSE
HALLMARK_INFLAMMATORY_RESPONSE
HALLMARK_APOPTOSIS
HALLMARK_KRAS_SIGNALING_UP
HALLMARK_APICAL_JUNCTION
HALLMARK_FATTY_ACID_METABOLISM
HALLMARK_ADIPOGENESIS
HALLMARK_ANDROGEN_RESPONSE
HALLMARK_OXIDATIVE_PHOSPHORYLATION
HALLMARK_PROTEIN_SECRETION
Pathway
Normalized Enrichment Score
−Log10(adj p−value)
B
Top 20 Pathways
Human Atrophic/Intermediate vs Luminal Clusters
HALLMARK_MYC_TARGETS_V1
HALLMARK_TNFA_SIGNALING_VIA_NFKB
DANG_MYC_TARGETS_UP
HALLMARK_INTERFERON_GAMMA_RESPONSE
HALLMARK_ALLOGRAFT_REJECTION
HALLMARK_MYC_TARGETS_V2
HALLMARK_P53_PATHWAY
HALLMARK_UNFOLDED_PROTEIN_RESPONSE
HALLMARK_INFLAMMATORY_RESPONSE
HALLMARK_E2F_TARGETS
HALLMARK_APOPTOSIS
HALLMARK_ESTROGEN_RESPONSE_EARLY
HALLMARK_EPITHELIAL_MESENCHYMAL_TRANSITION
HALLMARK_ESTROGEN_RESPONSE_LATE
HALLMARK_COMPLEMENT
HALLMARK_FATTY_ACID_METABOLISM
HALLMARK_ADIPOGENESIS
HALLMARK_PROTEIN_SECRETION
HALLMARK_ANDROGEN_RESPONSE
HALLMARK_OXIDATIVE_PHOSPHORYLATION
Pathway
Normalized Enrichment Score
−Log10(adj p−value)
C
Top 20 Pathways
Human Cancer vs Atrophic/Intermediate Clusters
HALLMARK_ANDROGEN_RESPONSE
HALLMARK_OXIDATIVE_PHOSPHORYLATION
HALLMARK_FATTY_ACID_METABOLISM
HALLMARK_ADIPOGENESIS
HALLMARK_TGF_BETA_SIGNALING
HALLMARK_HYPOXIA
HALLMARK_INFLAMMATORY_RESPONSE
HALLMARK_ALLOGRAFT_REJECTION
HALLMARK_MITOTIC_SPINDLE
HALLMARK_COMPLEMENT
HALLMARK_APOPTOSIS
HALLMARK_ESTROGEN_RESPONSE_EARLY
HALLMARK_IL6_JAK_STAT3_SIGNALING
HALLMARK_P53_PATHWAY
HALLMARK_PROTEIN_SECRETION
HALLMARK_UNFOLDED_PROTEIN_RESPONSE
HALLMARK_INTERFERON_ALPHA_RESPONSE
HALLMARK_MYC_TARGETS_V1
HALLMARK_INTERFERON_GAMMA_RESPONSE
HALLMARK_TNFA_SIGNALING_VIA_NFKB
Pathway
Normalized Enrichment Score
−Log10(adj p−value)
D
Human Epithelial Groups
Luminal
Atrophic/Intermediate
Basal
Cancer
UMAP_2
UMAP_1
E
MYC
UMAP_2
UMAP_1
F
MYC V1 Leading Edge
UMAP_2
UMAP_1
G
Human Tissue
H&E
PIN4
MYC
Normal
Adenocarcinoma
2x
16x
Basal
Cancer

**Figure S7**

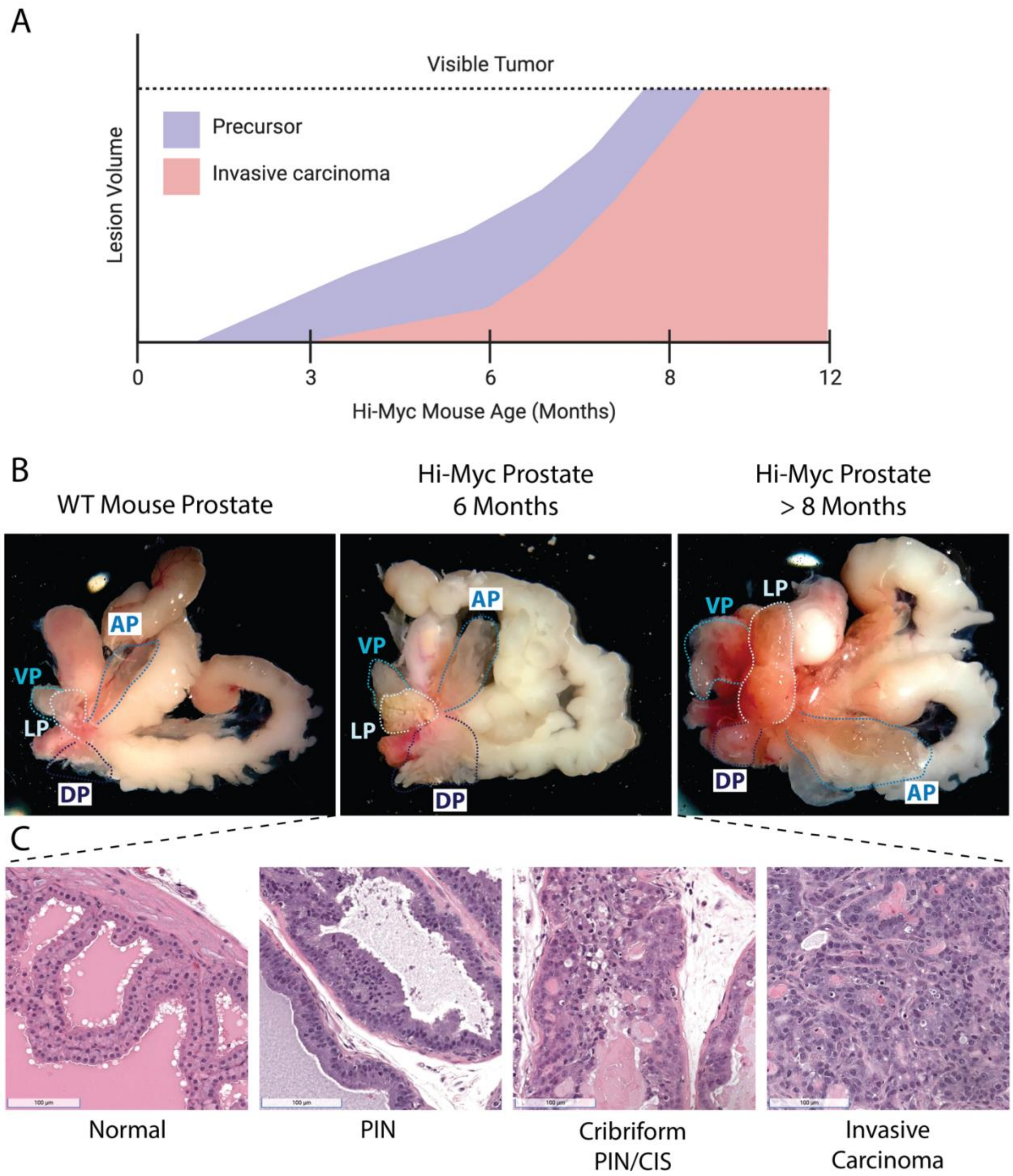
A
Visible Tumor
Precursor
Invasive carcinoma
Lesion Volume
0
3
6
8
12
Hi-Myc Mouse Age (Months)
B
WT Mouse Prostate
Hi-Myc Prostate
6 Months
Hi-Myc Prostate
> 8 Months
AP
VP
LP
DP
C
Normal
PIN
Cribriform
PIN/CIS
Invasive
Carcinoma

**Figure S8**

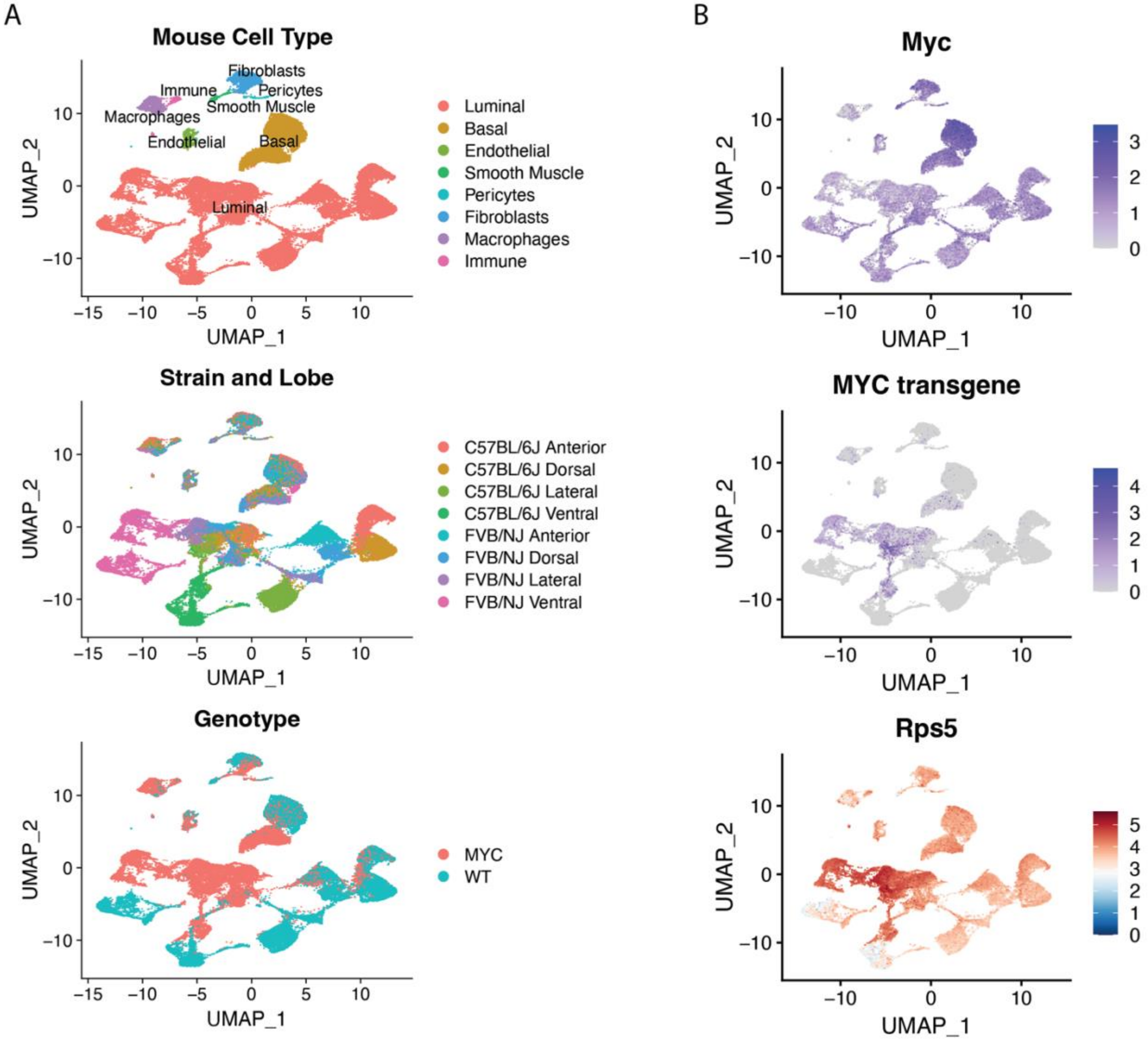
A
Mouse Cell Type
Fibroblasts
Immune
Pericytes
Smooth Muscle
Macrophages
Endothelial
Basal
Luminal
UMAP_2
UMAP_1
Luminal
Basal
Endothelial
Smooth Muscle
Pericytes
Fibroblasts
Macrophages
Immune
Strain and Lobe
C57BL/6J Anterior
C57BL/6J Dorsal
C57BL/6J Lateral
C57BL/6J Ventral
FVB/NJ Anterior
FVB/NJ Dorsal
FVB/NJ Lateral
FVB/NJ Ventral
Genotype
MYC
WT
B
Myc
MYC transgene
Rps5

**Figure S9**

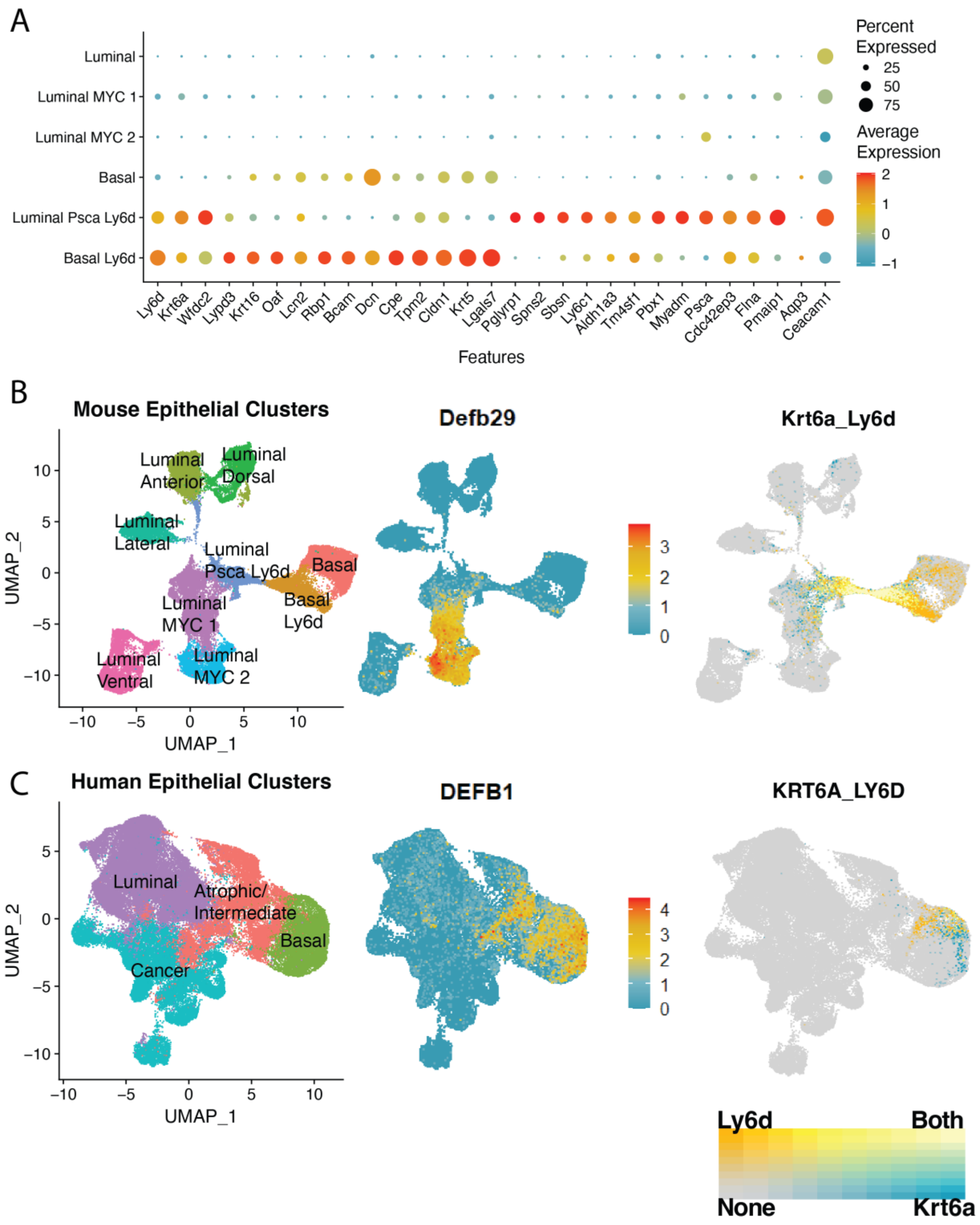

A
Percent Expressed
25
50
75
Average Expression
2
1
0
−1
Luminal
Luminal MYC 1
Luminal MYC 2
Basal
Luminal Psca Ly6d
Basal Ly6d
Ly6d
Krt6a
Wfdc2
Lypd3
Krt16
Oaf
Lcn2
Rbp1
Bcam
Dcn
Cpe
Tpm2
Cldn1
Krt5
Lgals7
Pglyrp1
Spns2
Sbsn
Ly6c1
Aldh1a3
Tm4sf1
Pbx1
Myadm
Psca
Cdc42ep3
Flna
Pmaip1
Aqp3
Ceacam1
Features
B
Mouse Epithelial Clusters
Luminal Anterior
Luminal Dorsal
Luminal Lateral
Luminal Psca Ly6d
Basal
Basal Ly6d
Luminal MYC 1
Luminal MYC 2
Luminal Ventral
UMAP_2
UMAP_1
Defb29
Krt6a_Ly6d
C
Human Epithelial Clusters
Luminal
Atrophic/ Intermediate
Basal
Cancer
UMAP_2
UMAP_1
DEFB1
KRT6A_LY6D
Ly6d
Both
None
Krt6a


**Figure S10**

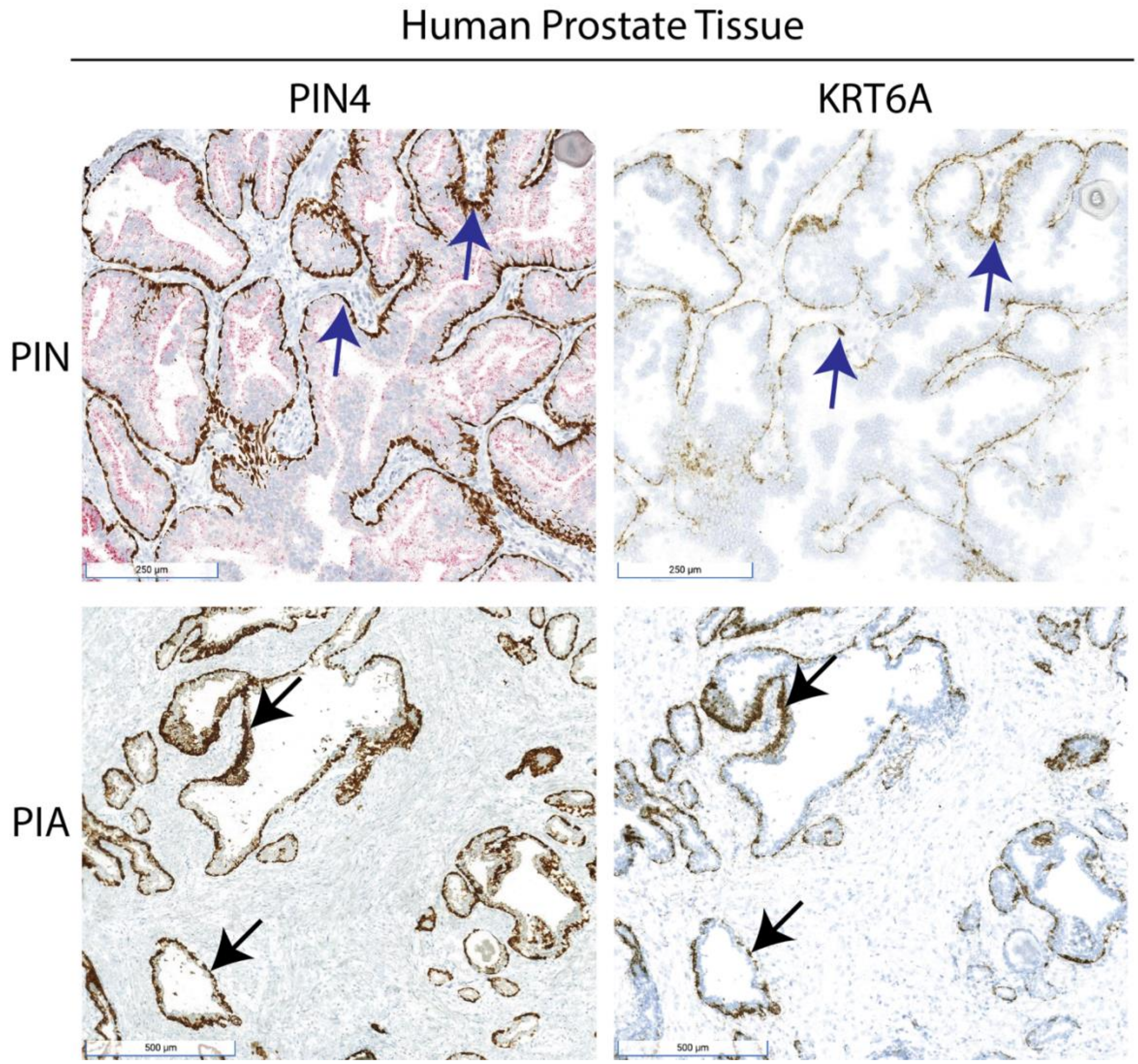
Human Prostate Tissue
PIN4
KRT6A
PIN
PIA
250 µm
250 µm
500 µm
500 µm

**Figure S11**

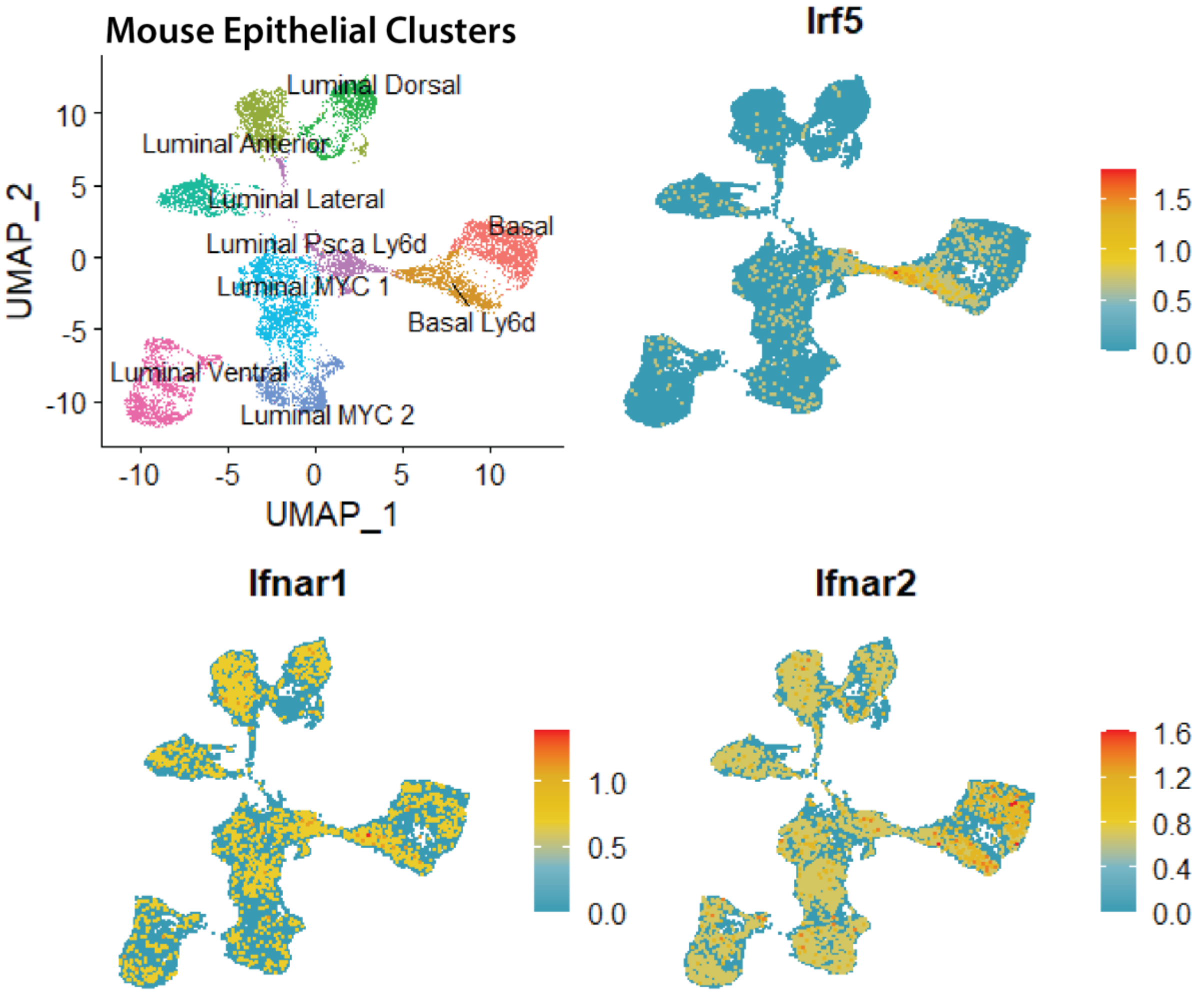
Mouse Epithelial Clusters
Luminal Dorsal
Luminal Anterior
Luminal Lateral
Luminal Psca Ly6d
Basal
Luminal MYC 1
Basal Ly6d
Luminal Ventral
Luminal MYC 2
UMAP_2
UMAP_1
Irf5
Ifnar1
Ifnar2

**Figure S12**

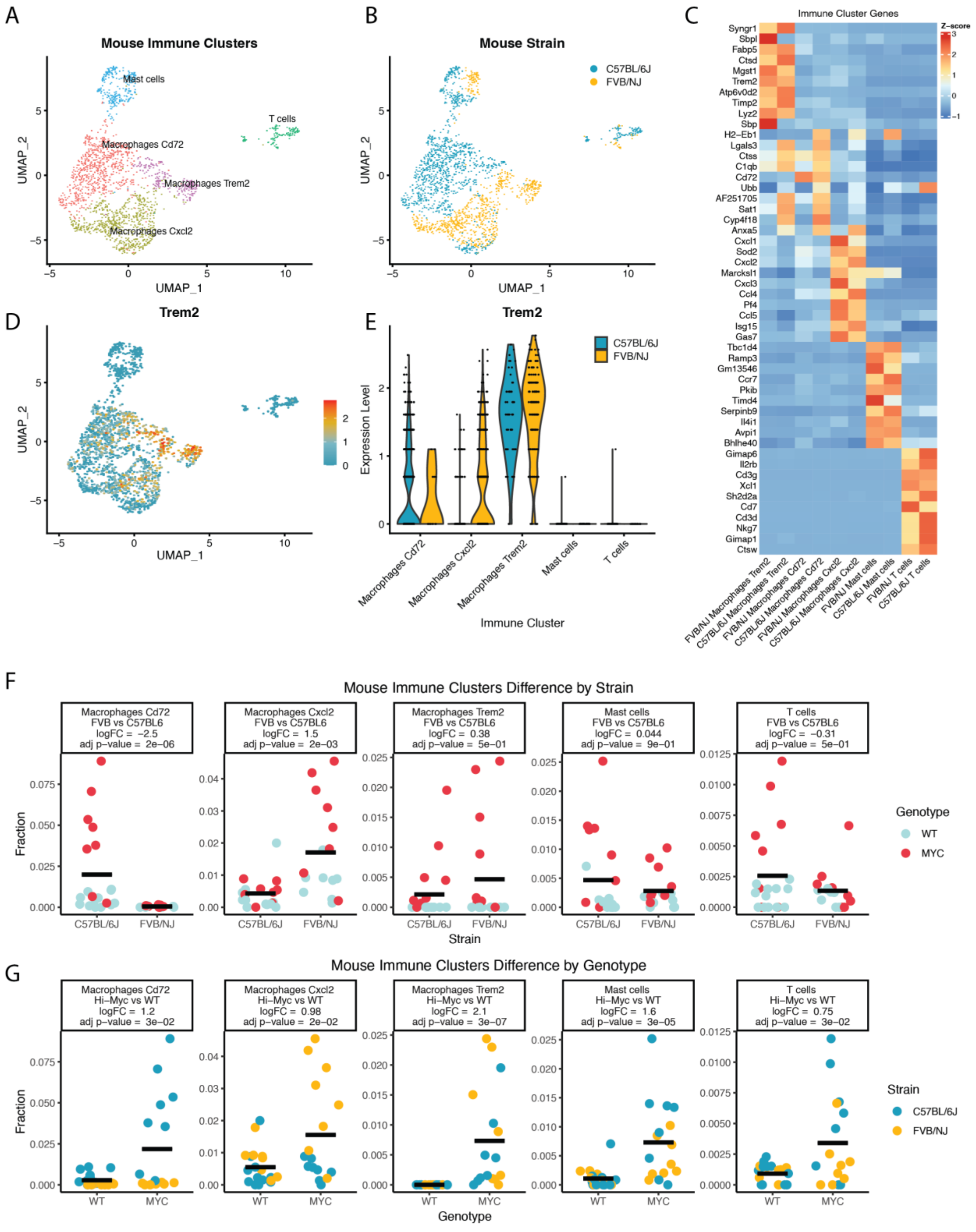
A
Mouse Immune Clusters
Mast cells
T cells
Macrophages Cd72
Macrophages Trem2
Macrophages Cxcl2
UMAP_1
UMAP_2
B
Mouse Strain
C57BL/6J
FVB/NJ
C
Immune Cluster Genes
Z-score
Syngr1
Sbpl
Fabp5
Ctsd
Mgst1
Trem2
Atp6v0d2
Timp2
Lyz2
Sbp
H2-Eb1
Lgals3
Ctss
C1qb
Cd72
Ubb
AF251705
Sat1
Cyp4f18
Anxa5
Cxcl1
Sod2
Cxcl2
Marcksl1
Cxcl3
Ccl4
Pf4
Ccl5
Isg15
Gas7
Tbc1d4
Ramp3
Gm13546
Ccr7
Pkib
Timd4
Serpinb9
Il4i1
Avpi1
Bhlhe40
Gimap6
Il2rb
Cd3g
Xcl1
Sh2d2a
Cd7
Cd3d
Nkg7
Gimap1
Ctsw
FVB/NJ Macrophages Trem2
C57BL/6J Macrophages Trem2
FVB/NJ Macrophages Cd72
C57BL/6J Macrophages Cd72
FVB/NJ Macrophages Cxcl2
C57BL/6J Macrophages Cxcl2
FVB/NJ Mast cells
C57BL/6J Mast cells
FVB/NJ T cells
C57BL/6J T cells
D
Trem2
E
Trem2
C57BL/6J
FVB/NJ
Expression Level
Macrophages Cd72
Macrophages Cxcl2
Macrophages Trem2
Mast cells
T cells
Immune Cluster
F
Mouse Immune Clusters Difference by Strain
Macrophages Cd72 FVB vs C57BL6 logFC = −2.5 adj p−value = 2e−06
Macrophages Cxcl2 FVB vs C57BL6 logFC = 1.5 adj p−value = 2e−03
Macrophages Trem2 FVB vs C57BL6 logFC = 0.38 adj p−value = 5e−01
Mast cells FVB vs C57BL6 logFC = 0.044 adj p−value = 9e−01
T cells FVB vs C57BL6 logFC = −0.31 adj p−value = 5e−01
Fraction
Strain
C57BL/6J
FVB/NJ
Genotype
WT
MYC
G
Mouse Immune Clusters Difference by Genotype
Macrophages Cd72 Hi−Myc vs WT logFC = 1.2 adj p−value = 3e−02
Macrophages Cxcl2 Hi−Myc vs WT logFC = 0.98 adj p−value = 2e−02
Macrophages Trem2 Hi−Myc vs WT logFC = 2.1 adj p−value = 3e−07
Mast cells Hi−Myc vs WT logFC = 1.6 adj p−value = 3e−05
T cells Hi−Myc vs WT logFC = 0.75 adj p−value = 3e−02
Fraction
Genotype
WT
MYC
Strain
C57BL/6J
FVB/NJ

**Figure S13**

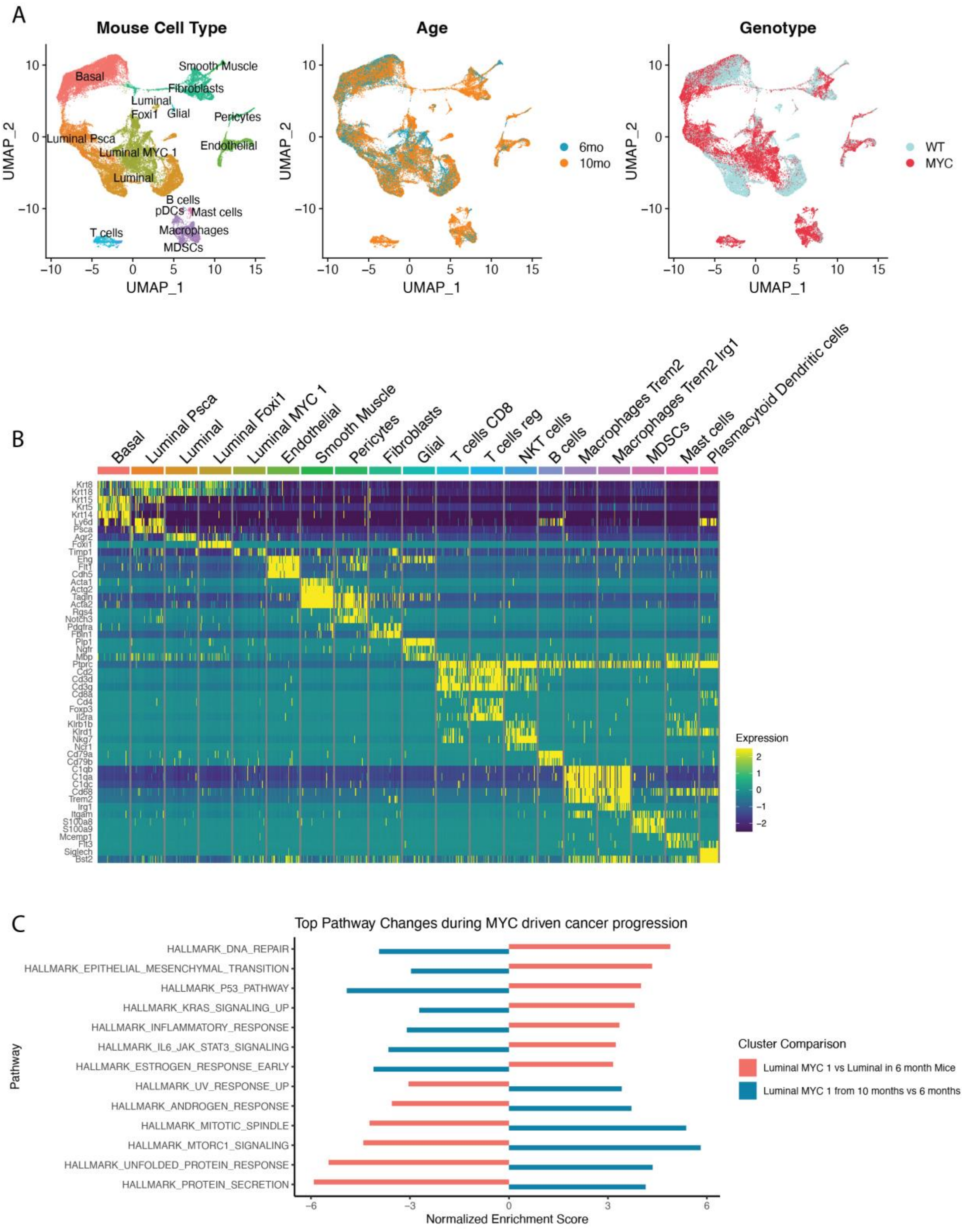
A
Mouse Cell Type
Age
Genotype
Basal
Smooth Muscle
Fibroblasts
Luminal
Foxi1
Glial
Pericytes
Luminal Psca
Endothelial
Luminal MYC 1
Luminal
B cells
pDCs
Mast cells
Macrophages
T cells
MDSCs
UMAP_1
UMAP_2
6mo
10mo
WT
MYC
B
Basal
Luminal Psca
Luminal
Luminal Foxi1
Luminal MYC 1
Endothelial
Smooth Muscle
Pericytes
Fibroblasts
Glial
T cells CD8
T cells reg
NKT cells
B cells
Macrophages Trem2
Macrophages Trem2 Irg1
MDSCs
Mast cells
Plasmacytoid Dendritic cells
Expression
C
Top Pathway Changes during MYC driven cancer progression
HALLMARK_DNA_REPAIR
HALLMARK_EPITHELIAL_MESENCHYMAL_TRANSITION
HALLMARK_P53_PATHWAY
HALLMARK_KRAS_SIGNALING_UP
HALLMARK_INFLAMMATORY_RESPONSE
HALLMARK_IL6_JAK_STAT3_SIGNALING
HALLMARK_ESTROGEN_RESPONSE_EARLY
HALLMARK_UV_RESPONSE_UP
HALLMARK_ANDROGEN_RESPONSE
HALLMARK_MITOTIC_SPINDLE
HALLMARK_MTORC1_SIGNALING
HALLMARK_UNFOLDED_PROTEIN_RESPONSE
HALLMARK_PROTEIN_SECRETION
Pathway
Normalized Enrichment Score
Cluster Comparison
Luminal MYC 1 vs Luminal in 6 month Mice
Luminal MYC 1 from 10 months vs 6 months

**Figure S14**

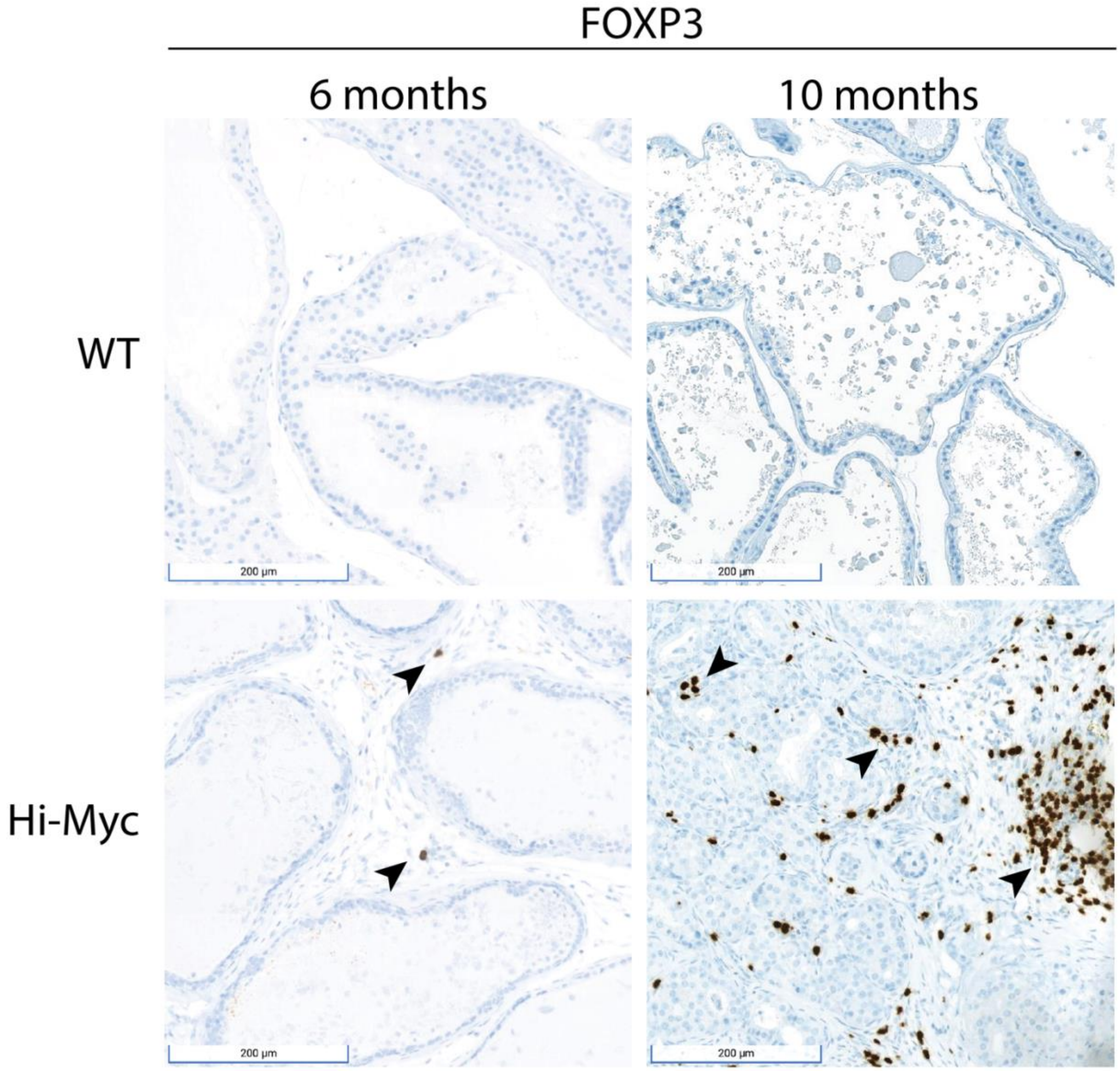


**Figure S15**

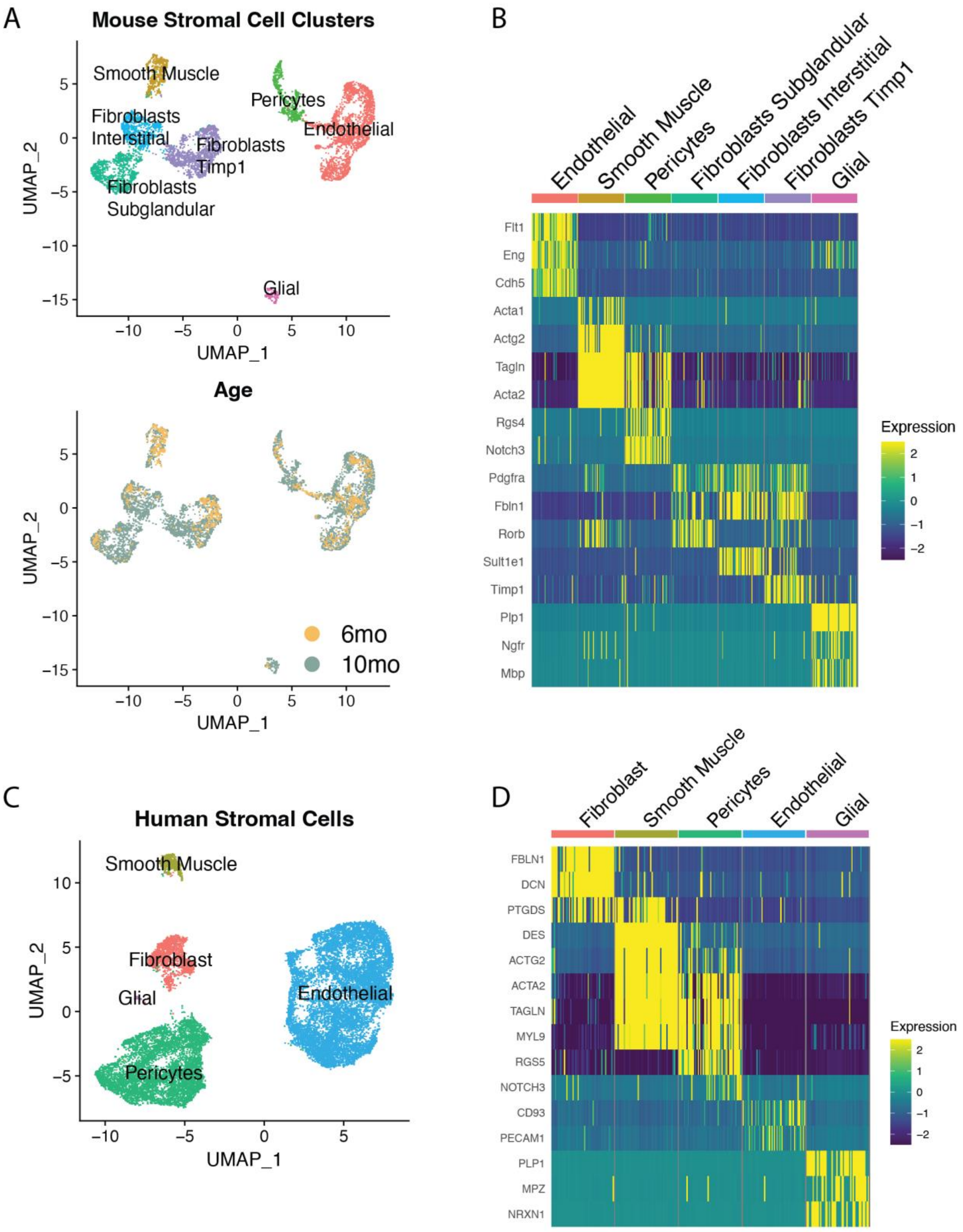
A
Mouse Stromal Cell Clusters
Smooth Muscle
Pericytes
Fibroblasts Interstitial
Endothelial
Fibroblasts Timp1
Fibroblasts Subglandular
Glial
UMAP_1
UMAP_2
Age
6mo
10mo
B
Endothelial
Smooth Muscle
Pericytes
Fibroblasts Subglandular
Fibroblasts Interstitial
Fibroblasts Timp1
Glial
Flt1
Eng
Cdh5
Acta1
Actg2
Tagln
Acta2
Rgs4
Notch3
Pdgfra
Fbln1
Rorb
Sult1e1
Timp1
Plp1
Ngfr
Mbp
Expression
C
Human Stromal Cells
Smooth Muscle
Fibroblast
Glial
Endothelial
Pericytes
D
Fibroblast
Smooth Muscle
Pericytes
Endothelial
Glial
FBLN1
DCN
PTGDS
DES
ACTG2
ACTA2
TAGLN
MYL9
RGS5
NOTCH3
CD93
PECAM1
PLP1
MPZ
NRXN1
Expression

**Figure S16**

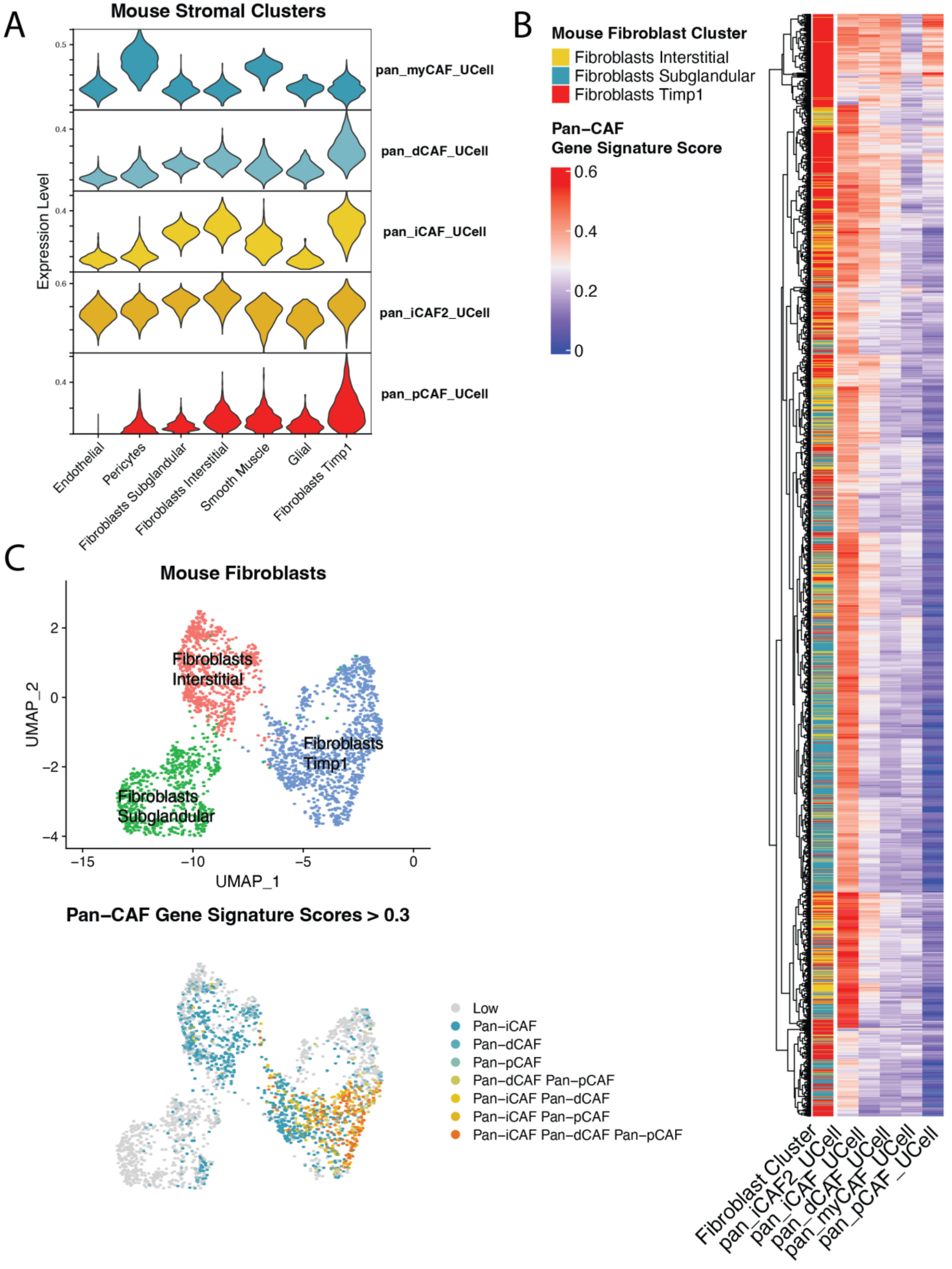
A
Mouse Stromal Clusters
Expression Level
pan_myCAF_UCell
pan_dCAF_UCell
pan_iCAF_UCell
pan_iCAF2_UCell
pan_pCAF_UCell
Endothelial
Pericytes
Fibroblasts Subglandular
Fibroblasts Interstitial
Smooth Muscle
Glial
Fibroblasts Timp1
B
Mouse Fibroblast Cluster
Fibroblasts Interstitial
Fibroblasts Subglandular
Fibroblasts Timp1
Pan−CAF
Gene Signature Score
0.6
0.4
0.2
0
Fibroblast Cluster
pan_iCAF2_UCell
pan_iCAF_UCell
pan_dCAF_UCell
pan_myCAF_UCell
pan_pCAF_UCell
C
Mouse Fibroblasts
UMAP_2
UMAP_1
Fibroblasts Interstitial
Fibroblasts Timp1
Fibroblasts Subglandular
Pan−CAF Gene Signature Scores > 0.3
Low
Pan−iCAF
Pan−dCAF
Pan−pCAF
Pan−dCAF Pan−pCAF
Pan−iCAF Pan−dCAF
Pan−iCAF Pan−pCAF
Pan−iCAF Pan−dCAF Pan−pCAF

**Figure S17**

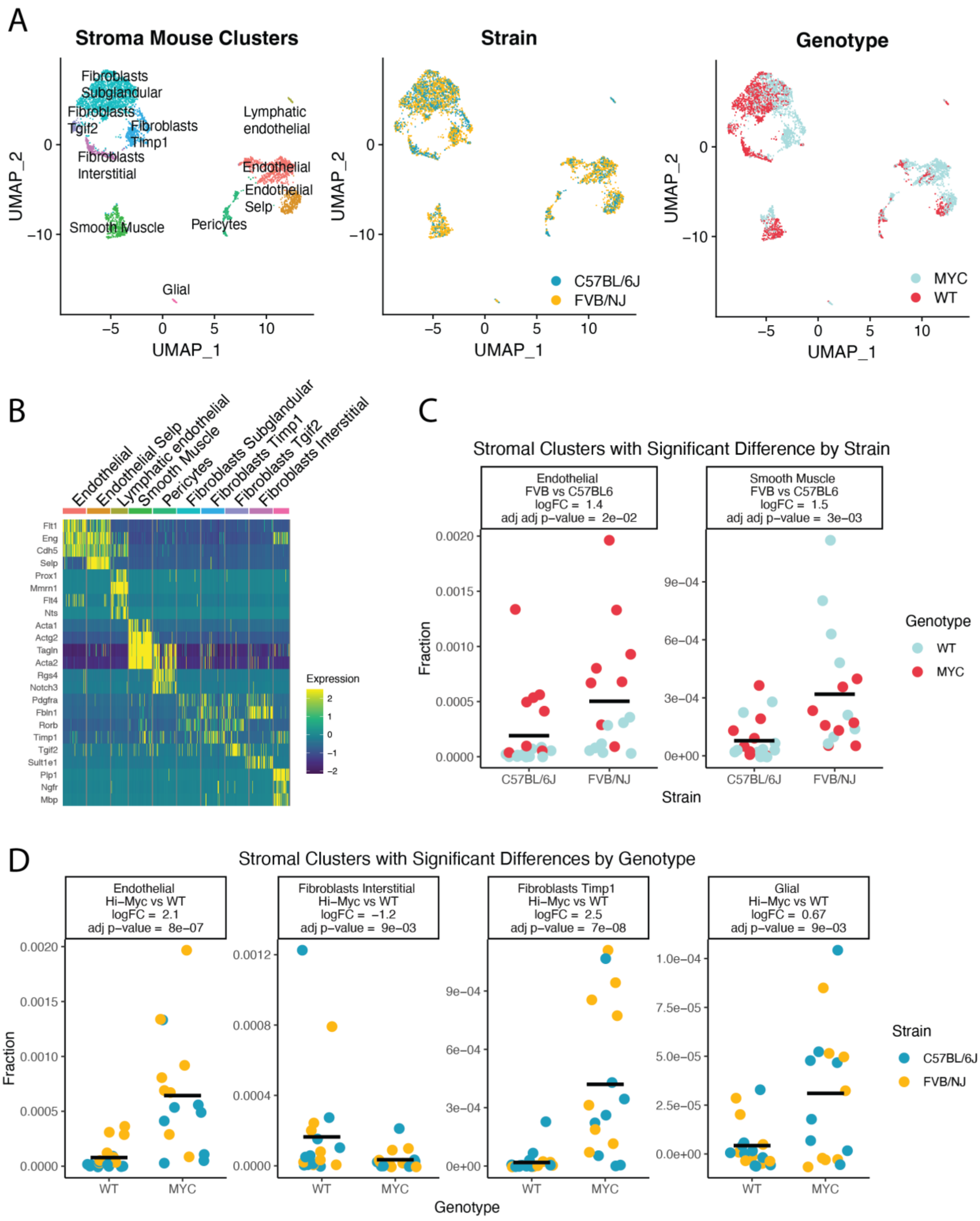
A
Stroma Mouse Clusters
Fibroblasts Subglandular
Fibroblasts Tgif2
Fibroblasts Timp1
Fibroblasts Interstitial
Lymphatic endothelial
Endothelial
Endothelial Selp
Smooth Muscle
Pericytes
Glial
UMAP_1
UMAP_2
Strain
C57BL/6J
FVB/NJ
Genotype
MYC
WT
B
Endothelial
Endothelial Selp
Lymphatic endothelial
Smooth Muscle
Pericytes
Fibroblasts Subglandular
Fibroblasts Timp1
Fibroblasts Tgif2
Fibroblasts Interstitial
Flt1
Eng
Cdh5
Selp
Prox1
Mmrn1
Flt4
Nts
Acta1
Actg2
Tagln
Acta2
Rgs4
Notch3
Pdgfra
Fbln1
Rorb
Timp1
Tgif2
Sult1e1
Plp1
Ngfr
Mbp
Expression
C
Stromal Clusters with Significant Difference by Strain
Endothelial
FVB vs C57BL6
logFC = 1.4
adj adj p−value = 2e−02
Smooth Muscle
FVB vs C57BL6
logFC = 1.5
adj adj p−value = 3e−03
Fraction
Strain
C57BL/6J
FVB/NJ
Genotype
WT
MYC
D
Stromal Clusters with Significant Differences by Genotype
Endothelial
Hi−Myc vs WT
logFC = 2.1
adj p−value = 8e−07
Fibroblasts Interstitial
Hi−Myc vs WT
logFC = −1.2
adj p−value = 9e−03
Fibroblasts Timp1
Hi−Myc vs WT
logFC = 2.5
adj p−value = 7e−08
Glial
Hi−Myc vs WT
logFC = 0.67
adj p−value = 9e−03
Fraction
Genotype
WT
MYC
Strain
C57BL/6J
FVB/NJ

**Figure S18**
